# Immunogenicity of non-canonical HLA-I tumor ligands identified through proteogenomics

**DOI:** 10.1101/2022.11.07.514886

**Authors:** Maria Lozano-Rabella, Andrea Garcia-Garijo, Jara Palomero, Anna Yuste-Estevanez, Florian Erhard, Juan Martín-Liberal, Maria Ochoa de Olza, Ignacio Matos, Jared J. Gartner, Michael Ghosh, Francesc Canals, August Vidal, Josep Maria Piulats, Xavier Matias-Guiu, Irene Braña, Eva Muñoz-Couselo, Elena Garralda, Andreas Schlosser, Alena Gros

## Abstract

Tumor antigens are central to antitumor immunity. Recent evidence suggests that peptides from non-canonical (nonC) aberrantly translated proteins can be presented on HLA-I by tumor cells. Here, we investigated the immunogenicity of nonC tumor HLA-I ligands (nonC-TL) to better understand their contribution to cancer immunosurveillance and their therapeutic applicability. Using proteogenomics, we identified 517 nonC-TL from 9 patients with melanoma, gynecological, and head and neck cancer. We found no recognition of the 507 nonC-TL tested by autologous *ex vivo* expanded tumor reactive T-cell cultures while the same cultures demonstrated reactivity to mutated, cancer-germline, or melanocyte differentiation antigens. However, *in vitro* sensitization of donor peripheral blood lymphocytes against 170 selected nonC-TL, led to the identification of T-cell receptors (TCRs) specific to three nonC-TL, two of which mapped to the 5’ UTR regions of HOXC13 and ZKSCAN1, and one mapping to a non-coding spliced variant of C5orf22C. T cells targeting these nonC-TL recognized cancer cell lines naturally presenting their corresponding antigens. Expression of the three immunogenic nonC-TL was shared across tumor types and barely or not detected in normal cells. Our findings predict a limited contribution of nonC-TL to cancer immunosurveillance but demonstrate they may be attractive novel targets for widely applicable immunotherapies.

## Introduction

Tumor antigens are central to antitumor immunity. Peptides derived from tumor antigens presented on HLA molecules (pHLA) on the surface of cancer cells can elicit protective and therapeutic T-cell responses(1). The existence of T cells targeting non-mutated tumor-associated antigens (TAA) and cancer-germline antigens (CGA) in cancer patients is well established(2,3). Their shared expression in a substantial fraction of tumors has led to the development of widely applicable vaccines or T cell-based therapies (4,5). However, off-tumor toxicities have been reported(6–8). Technological advances in next-generation sequencing (NGS) and tandem mass spectrometry coupled with high-throughput immunological and HLA multimer screens have expedited the systematic discovery of the personalized landscape of antigens contributing to tumor immunogenicity. Accumulating evidence demonstrates that neoantigens arising from non-synonymous somatic mutations (NSM) greatly contribute to the immunogenicity of human tumors. For instance, neoantigen-specific T cells are frequently detected in cancer patients (9–13) and mutational load correlates with the clinical benefit of immune checkpoint blockade (ICB)(14). Their foreign nature and high tumor specificity together with the antitumor responses observed following transfer of neoantigen-specific T cells in selected patients (15–18) render these attractive targets. Yet, existing techniques still fail to capture most antigens targeted by tumor-reactive T cells and this constitute a major obstacle for the development of immunotherapy.

Tumor antigen discovery efforts thus far have largely investigated the immunogenicity of selected genomically annotated proteins or NSM in coding regions, limited to only 2% of the genome. However, up to 75% of the genome can be transcribed and, potentially, translated(19). Emerging data demonstrate that peptides derived from alternative open reading frames (ORF) or from allegedly non-coding regions referred to as non-canonical (nonC) or cryptic antigens are frequently presented on HLA-I molecules (20–23). A fraction of such aberrant translation events has been postulated to be specifically presented on tumor cells, thus substantially expanding the repertoire of targetable tumor antigens (24–27). Their non-mutated nature and occasional shared presentation across different tumors, has further attracted attention to nonC proteins as targets for immunotherapy.

Despite the potential of tumor-specific nonC HLA-I ligands as a source of tumor antigens, their systematic identification in humans remains challenging. Spontaneous T-cell responses against peptides derived from nonC proteins have been rarely identified using cumbersome and time-consuming immunological screens of tumor cDNA libraries (28–30). A growing number of recent studies have exploited immunopeptidomics to identify these antigens (22–27,31), but their immunogenicity and their selective expression in cancer remains largely unexplored. Here, we investigated the presentation and immunogenicity of nonC antigens across different cancer types to better understand their contribution to cancer immunosurveillance and to address their therapeutic potential. We employed a proteogenomics pipeline(20) to identify nonC HLA-I ligands derived from off-frame translation of coding sequences and non-coding regions (UTR, ncRNA, intronic and intergenic) in patient-derived tumor cell lines (TCLs) of different histological types. We further modified the pipeline to select peptides preferentially presented by cancer cells and evaluated their natural or induced immunogenicity by assessing pre-existing and *in vitro*-sensitized T-cell responses.

## Results

### Non-canonical tumor HLA-I ligands are frequently identified in patient-derived tumor cell lines

We first sought to determine whether tumor-specific nonC HLA-I ligands could be identified in 9 short-term cultured TCL derived from four gynecological cancer (Gyn), three melanoma (Mel) and two head and neck (H&N) cancer patients (Supplemental Table 1). These samples were selected irrespective of the tumor histology, solely based on the availability of matched *ex vivo* expanded TIL and/or peripheral blood tumor-reactive lymphocyte populations.

To this end, peptides bound to HLA-I were isolated and analyzed by liquid chromatography coupled to tandem mass spectrometry (LC-MS/MS) using state-of-the-art procedures. Amino acid (Aa) sequences were identified through a previously described pipeline, Peptide-PRISM(20), with some modifications (Figure 1A). Briefly, for each MS spectrum, the top 10 candidates were first identified by *de novo* sequencing and later mapped to a database including the 3-frame transcriptome and 6-frame genome. Additionally, whole-exome sequencing (WES) information of each TCL was included to interrogate the presentation of mutated peptides derived from cancer-specific NSM. The false-discovery rate (FDR) was calculated independently for each category considering the search space and peptide length in a stratified mixture model as previously described (20). Following this strategy and selecting a 1% FDR, we identified 839 nonC peptides presented on HLA-I in all the TCLs studied, ranging from 0.5% to 5.4% of the total eluted peptides (Figure 1B).

**Figure 1.**
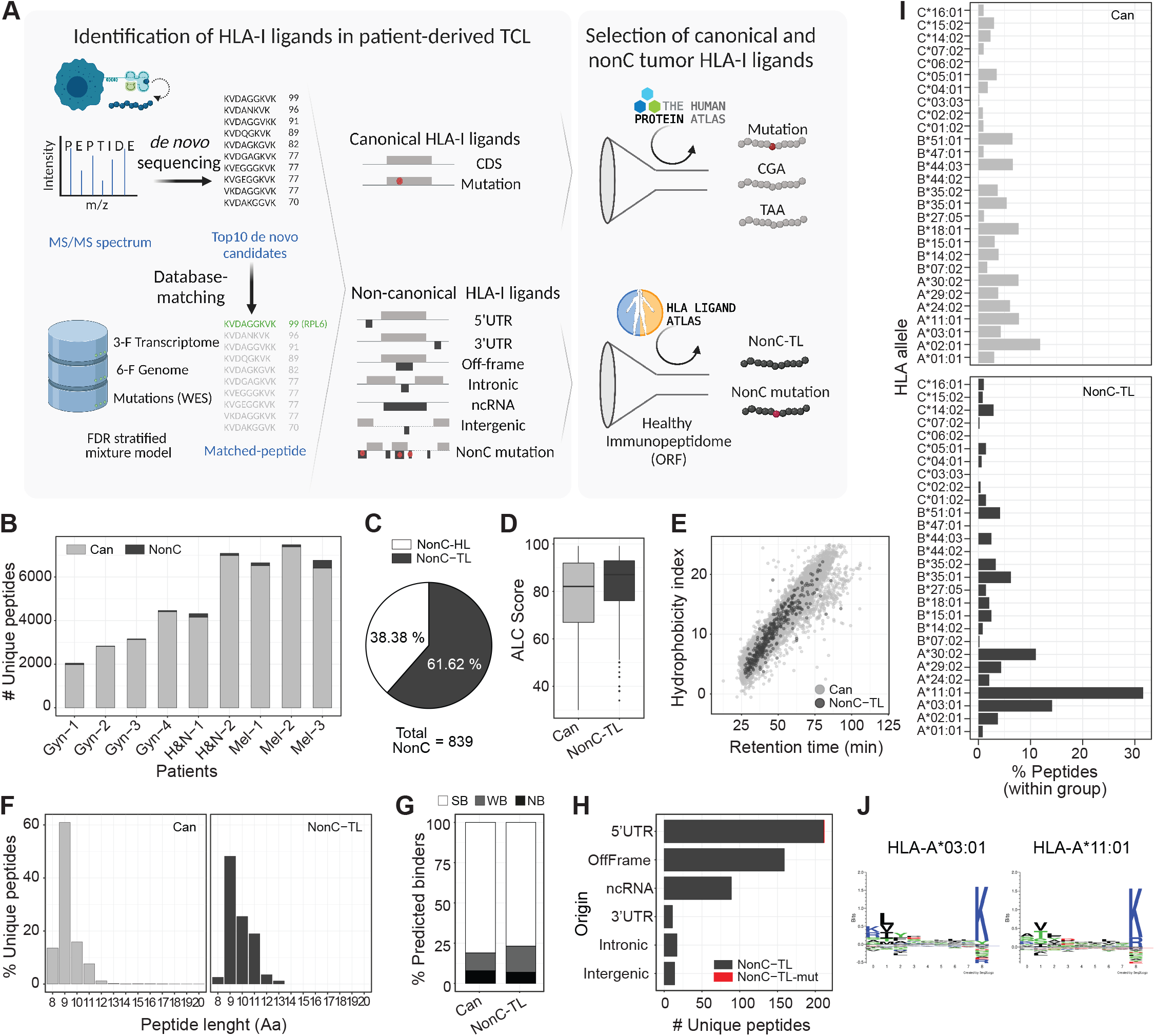
Identification and characteristics of non-canonical HLA-I ligands presented by patient-derived tumor cell lines. (**A**) Diagram depicting the pipeline used to identify HLA-I ligands derived from canonical (Can) and non-canonical (nonC) proteins presented by patient-derived TCL. The Top10 candidates for each MS spectrum were identified by de novo sequencing and aligned to a database containing the 3-frame transcriptome, 6-frame genome and the NSM identified by WES. The FDR was calculated for each category shown using a stratified mixture model (left panel). All canonical peptides containing mutations as well as peptides derived from CGA or TAA were further studied. For nonC HLA-I ligands, healthy immunopeptidomics data from HLA ligand atlas was used to filter out peptides presented in healthy tissues at the ORF level to obtain the nonC-TL (right panel). Image created with BioRender. (**B**) Number of canonical and non-canonical HLA-I peptides identified per patient. (**C**) Percentage of nonC peptides derived from predicted ORF present or absent in a healthy tissue immunopeptidome data set (nonC-HL and nonC-TL, respectively). (**D**) ALC identification score of canonical and nonC-TL. (**E**) Predicted hydrophobicity index (y axis) and retention time (x axis). Each dot represents a unique peptide sequence. (**F**) Length distribution of unique HLA-I peptide sequences. Only peptides <20 Aa are depicted. (**G**) Percentage of peptides predicted to bind to the patient specific HLA alleles according to NetMHCPan4.0. Peptides were categorized into strong binders (SB; %-tile ran ≤ 0.5), weak binders (WB; %-tile rank =0.5-2) or non-binders (NB; %-tile rank > 2). (**H**) Number of nonC-TL originated from each of the ORF categories noted. (**I**) HLA allele binding preference of Can and nonC-TL. For each peptide, only the min rank predicted by NetMHCpan4.0 was considered. (**J**) Consensus peptide binding motif of the two HLAs predicted to present the majority of the nonC peptides identified. Image downloaded from NetMHCpan4.0 motif viewer. In all the analyses shown, the FDR threshold was set at 0.01 and ALC score at 30.

In order to select nonC peptides preferentially presented by tumor cells, immunopeptidomics data from samples available from the HLA ligand atlas (32) was used to filter out peptides known to be presented in healthy tissues (Figure 1A). Although LC-MS/MS is less sensitive than RNA-seq, this technique is capable of capturing canonical as well as aberrant translation. Given that we leveraged an immunopeptidomics database derived from donors presenting a fraction of all potential HLA alleles, nonC peptides were excluded at the ORF level rather than the Aa sequence to overcome a potential bias toward frequent alleles. As a result, we found that from a total of 839 unique nonC peptides detected in our tumor samples, 322 (38.38%) were predicted to derive from ORFs also present in healthy tissue (nonC-HL). Hence, 517 (61.6%) were considered preferentially presented on tumor HLA-I and referred to as non-canonical tumor ligands (nonC-TL) (Figure 1C). NonC-TL displayed similar characteristics as peptides derived from canonical proteins such as the MS identification score (ALC) or the correlation of the retention time with the hydrophobicity index (Figure 1D and 1E). Additionally, nonC-TL exhibited expected HLA-I ligand features as shown by the length distribution ranging from 8-12 Aa and the high percentage of peptides predicted to bind to the patient’s HLA alleles according to NetMHCpan4.0 (Figure 1F and 1G). Moreover, as with the HLA-I peptides derived from canonical proteins, most of the nonC-TL were validated by MS using synthetic or isotope labeled peptides, (Supplemental Figure 1-2). Altogether, these analyses indicate that our approach accurately identified the HLA-I ligand repertoire including nonC-TL presented by patient-derived TCL.

Next, we evaluated the genomic origin of the identified nonC-TL. Consistent with previous studies (20,33,34), we found that translation of 5’UTR was the main origin followed by off-frame and non-coding RNA (ncRNA) (Figure 1H). Peptides derived from 3’UTR, intronic and intergenic regions were less frequently detected. In addition, one nonC peptide derived from a 5’UTR containing a tumor-specific mutation was detected in patient Gyn-3 (Figure 1H). We noticed a clear bias in the HLA-I binding preference distribution of nonC ligands towards HLA-A*11:01 and HLA-A*03:01 alleles according to NetMHCpan4.0 (Figure 1I and Supplemental Figure 3). Both HLA alleles bind peptides with a similar motif containing basic residues at p9 (Figure 1J), a unique feature among all the alleles studied (Supplemental Figure 4). Indeed, the binding preference of nonC peptides to both HLA-A*11:01 and HLA-A*03:01 alleles has been previously reported in other immunopeptidomics studies (20,21), however the exact mechanism underpinning this bias is still unknown. Overall, our results showed that nonC-TL are frequently detected in patient-derived TCLs. Importantly, 76 of our nonC-TL were found in published HLA-I immunopeptidomics datasets from melanoma resection samples (35,36) (Supplemental Figure 5), supporting that HLA-I presentation of nonC-TL is not an artifact of *in vitro* cultured cells and that these peptides can be naturally presented *in vivo*.

### NonC-TL constitute an abundant source of candidate tumor antigens

To examine whether nonC-TL represent an attractive source of tumor antigens that could be exploited therapeutically we first compared the number of nonC-TL eluted from HLA-I to those derived from conventional tumor antigen sources, including peptides encoded by canonical coding regions derived from somatically mutated gene products, and from TAA such as CGA or melanoma-associated antigens arising from melanocyte differentiation proteins. While the number of HLA-I ligands derived from canonical tumor antigens ranged from 24 to 36 peptides, nonC-TL outnumbered the other categories, with 517 unique tumor antigen candidates in all the TCL studied (Figure 2A). Of note, this observation was consistent across most of the patients studied (Figure 2B). In more detail, a total of 33 mutated peptides were detected in 6 out of 9 patients. Despite the number of eluted HLA-I ligands containing mutations was low compared to the total NSM identified by WES, these results are in line with previous immunopeptidomics studies where few mutations are typically detected(36–38) (Figure 2B and 2C). Moreover, 36 peptides derived from 12 genes encoding for CGA and 24 peptides derived from 5 melanoma-associated antigens were identified in 8 out of 9 patients (Figure 2D). Furthermore, nonC-TL mainly originated from 5’UTR and off-frame translation in most patients, while peptides derived from intergenic and intronic regions were not or barely detected (Figure 2E).

**Figure 2.**
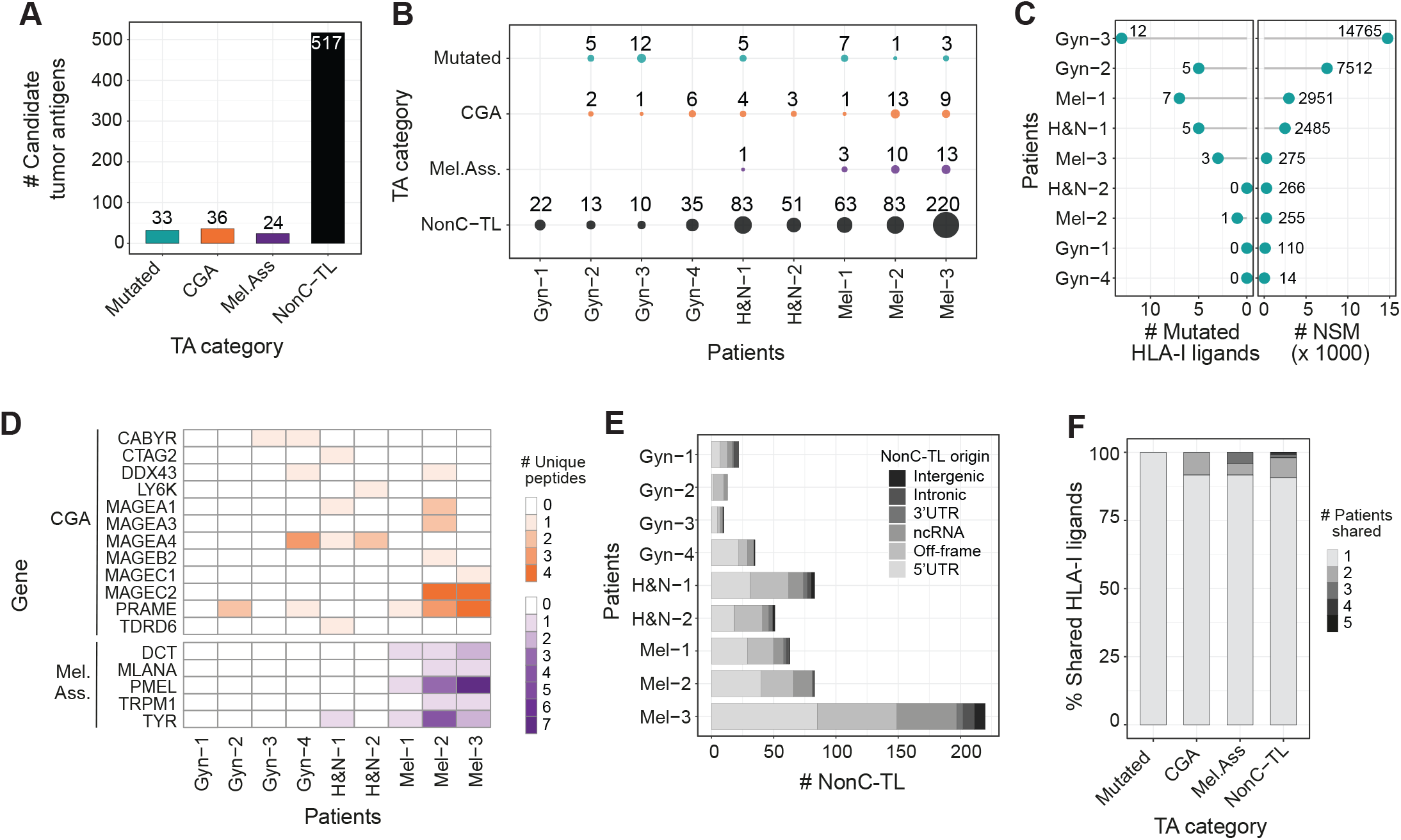
Candidate tumor antigens presented on HLA-I from patient-derived TCL identified through proteogenomics. (**A**)Total number of unique peptide sequences derived from candidate tumor antigens (TA) by category. Data from all patients was pooled together. Only unique peptide sequences were considered. (**B**) The number and category of TA are displayed for each TCL. (**C**) The number of mutated peptides eluted from HLA-I (left) and number of NSM identified by WES (right) are displayed for each patient. (**D**) Heat map displaying the number of epitopes derived from specific CGA or melanoma-associated antigens per patient. (**E**) Number of nonC-TL originated from each ORF category per patient. (**F**) Percentage of candidate TA uniquely identified in one patient or shared by TA category. The FDR threshold was set at 0.02 for mutations and 0.01 for all other categories.

One advantage of exploiting non-mutated antigens over private mutations as targetable tumor antigens is the fact that they can be shared across patients, which could facilitate the development of off-the-shelf vaccines or T-cell-based therapies. Similar to CGA or melanoma-associated antigens, we observed that ∼10% of nonC-TL were shared among at least 2 patients (Figure 2F). In contrast, all mutated HLA-I ligands detected were derived from private mutations and thus, exclusively identified in a single patient. Of note, nonC-TL showed the highest number of shared peptides, including one and three sequences identified in 5 and 4 patients, respectively (Figure 2F). Altogether, this data highlights nonC-TL as promising targets for the development of therapeutic interventions since they constitute a broader spectrum of candidate tumor antigens compared to peptides derived from mutations, CGA, or melanoma-associated antigens and can be shared across patients.

### Tumor-reactive T cells in cancer patients preferentially recognize neoantigens and TAA rather than nonC-TL

To further evaluate the role of nonC-TL in cancer immunosurveillance we assessed the presence of spontaneous T-cell responses targeting personalized candidate tumor antigens including nonC-TL, mutated, CGA and melanoma-associated antigens in the nine cancer patients. To this end, *ex vivo* expanded tumorinfiltrating (TIL) or peripheral blood lymphocyte (PBL) populations reactive to their corresponding autologous TCL were co-cultured with autologous antigen presenting cells (APC) pulsed with the synthetic peptides encoding the candidate antigens identified through proteogenomics. T-cell reactivity was tested by IFN-γ release and the upregulation of the activation cell-surface marker 4-1BB assessed by ELISPOT and flow cytometry, respectively (Figure 3A, 3B, and Supplemental Figure 6).

In patient Mel-3, we detected T-cell reactivity to at least one candidate peptide in 5 out of 11 different tumor-reactive TIL populations interrogated. TILs recognized two neoantigens (ETV1_p.E455K_ and GEMIN5_p.S1360L_) and two immunogenic peptides derived from melanoma-associated antigens (PMEL and MLANA). However, no reactivity was detected to any of the 9 peptides derived from CGA nor the 215 nonC-TL tested (Figure 3A). To further confirm and characterize the T-cell responses observed, antigen-specific T cells were enriched by flow cytometry based-sorting of 4-1BB^+^ cells followed by *ex vivo* expansion (Figure 3C). The neoantigen-specific enriched populations showed a higher response to the mutated peptide than to the wild type (Wt) counterparts as shown by peptide titration experiments, although T cells exhibited a variable functional avidity to their cognate antigen (Figure 3D). ETV1_p.E455K_ and GEMIN5_p.S1360L_ were restricted to HLA-B*35:01 and HLA-A*11:01, respectively (Figure 3E). Importantly, co-culture experiments showed that both neoantigen-specific T cells isolated recognized the autologous TCL, ultimately demonstrating that these peptides are naturally processed and presented by the tumor (Figure 3C and 3E).

**Figure 3.**
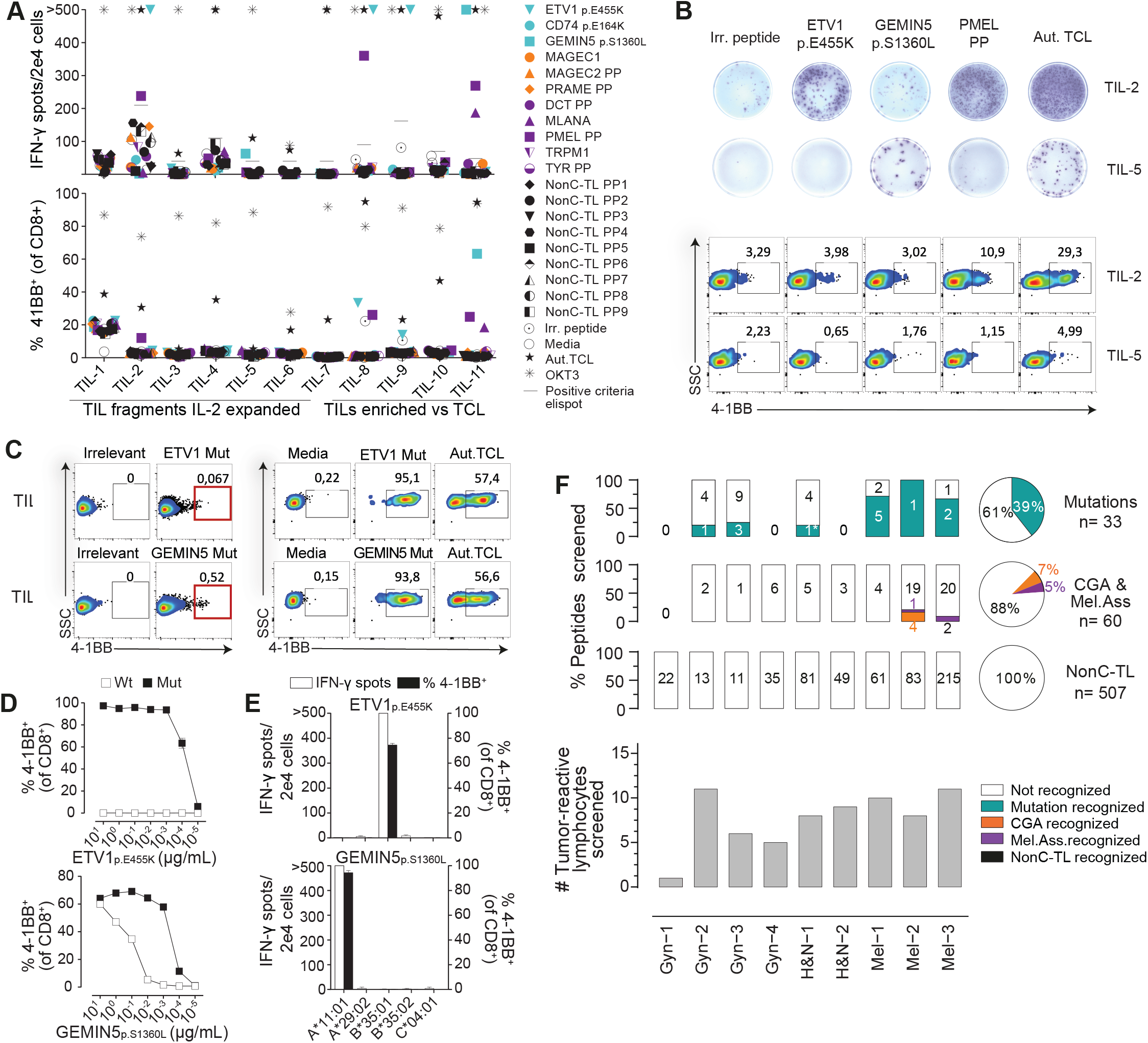
Pre-existing T-cell responses to candidate tumor antigens in cancer patients. For each patient, reactivity was evaluated by co-incubating 2e4 T cells (TIL o PBL sorted based on specific markers, e.g., PD1^hi^), with 2e5 autologous APC pulsed with 1 μg/mL of selected peptides either alone or in pools (PP). IFN-γ ELISPOT and 4-1BB upregulation by FACS were used to measure T-cell responses after 20 h. (**A**) Reactivity to tumor antigen candidates for Mel-3. The number of IFN-γ spots per well (top panel) and the percentage of cells expressing 4-1BB (bottom panel) are shown. Mutated peptides are plotted in turquoise, CGA in orange, melanoma-associated in purple, and nonC-TL in black. PP, peptide pool. (**B**) Representative ELISPOT results (top) and flow cytometry plots (bottom) for TIL-2 and TIL-5 from Mel-3 with the targets specified. (**C**) TIL populations recognizing the mutated HLA-I peptides indicated were enriched by flow cytometry sorting of 4-1BB^+^ lymphocytes and expanded for 14 days. Plot showing gates used for sorting (left) and recognition of the targets specified after expansion (right). (**D**) T cell reactivity of neoantigen-enriched T-cell populations to serial dilutions of the wild type (Wt) or mutant (Mut) ETV1_p.E455K_ and GEMIN5_p.S1360L_ peptides. (**E**) Neoantigen-enriched populationwere co-cultured with COS-7 cells transfected with the indicated individual HLA-I alleles and pulsed with the corresponding peptides to determine the restriction element. (**F**) Summary of the reactivity against candidate tumor antigens in all patients studied. The percentage and the absolute number of recognized and non-recognized peptides within each category are shown per patient (bar plot) and for all the patients studied (pie chart). The number of tumor-reactive lymphocytes tested for each patient is shown on the bottom. Plotted cells were gated on live CD3^+^CD8^+^lymphocytes. ‘>‘ denotes greater than 500 spots/2e4 cells. *Mutation recognized previously identified. Experiments were performed twice.

We used the same strategy to identify and characterize pre-existing T-cell responses targeting candidate tumor antigens from all patients included in the study (Figure 3F). In total, screening of preexisting tumor-reactive T cells for recognition of 600 tumor antigen candidates led to the detection of 20 immunogenic peptides in the 9 cancer patients studied (Figure 3F; Table 1). Of note, we could detect T-cell responses to at least one neoantigen in the six patients in which neoantigen candidates were detected through immunopeptidomics. Overall, 13 out of the 33 mutated HLA-I ligands tested were immunogenic, representing 39% of the total neoantigen candidates. Furthermore, four immunogenic peptides derived from CGA and three derived from melanoma-associated antigens were also recognized by naturally occurring tumor-reactive T cells. A detailed characterization of the antigen-specific T cells isolated including autologous tumor recognition, HLA restriction, functional avidity and Wt counterpart recognition for neoantigens is shown in Supplemental Figure 7-10. In contrast, none of the 507 unique nonC-TL candidates interrogated were able to elicit a recall immune response in any of the studied patients. Altogether, these results reveal that although nonC-TL were frequently detected in TCL, antigens derived from canonical regions were preferentially recognized by tumor-reactive lymphocytes.

**Table 1.**
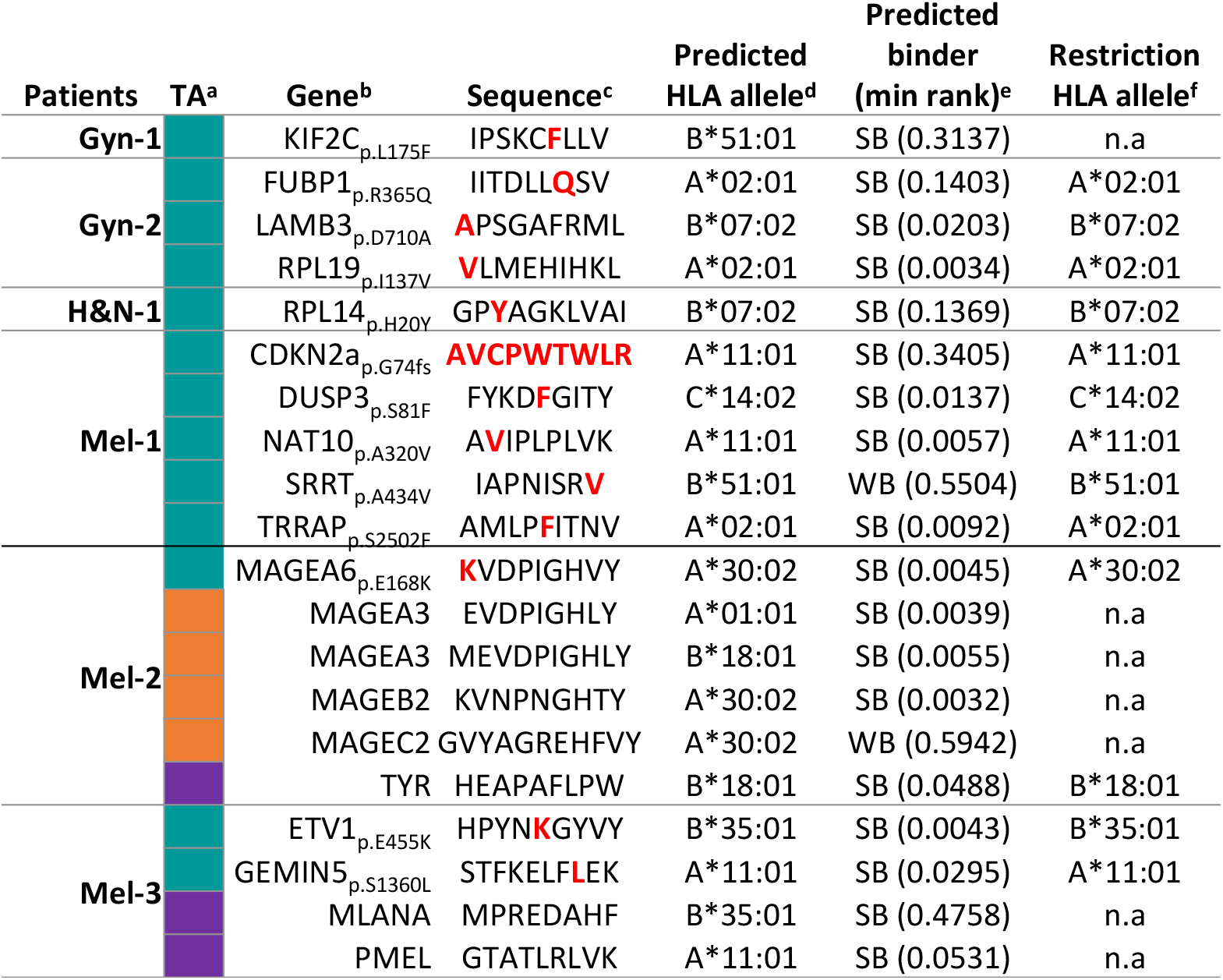
Immunogenic tumor antigens identified in cancer patients. ^a^The category of the tumor antigen (TA) is depicted in colors; mutated in turquoise, cancer-germline in orange and melanoma-associated in purple. ^b^Gene symbol, the amino acid change and position in the protein are shown. ^c^Mutated amino acids are highlighted in red letters. ^d,e^HLA predicted binding affinity using NetMHCpan4.0, only the allele with the minimal rank is shown. SB: Strong binders (%-tile rank ≤2); NB: non-binders (%-tile rank >2). ^f^Restriction element was evaluated experimentally by co-incubating reactive lymphocytes with COS-7 cells transfected with plasmids encoding for the individual HLA followed by peptide pulsing. T-cell responses were measured by IFNγ elispot and 4-1BB upregulation. n.a=non-assessed

### NonC-TL recognized by *in vitro* sensitized T cells are shared across patient-derived TCL

Although we did not detect pre-existing T-cell responses targeting nonC-TL, we reasoned that these antigens could still be immunogenic, and thus could represent attractive targets for T-cell therapies or vaccines. To address this question, we investigated the presence of naïve T cells specific to nonC-TL-in the repertoire of healthy individuals carrying the corresponding HLA restriction alleles. As such peptide specific TCR are in very low frequency we sought to enrich nonC-TL-specific T cells through *in vitro* sensitization (IVS) in a non-autologous HLA-matched setting (Figure 4A).

Out of the total 507 nonC-TL identified in all patients, we selected 170 peptides predicted to bind to HLA-A*11:01 according to NetMHCpan4.0, an allele expressed in 14% of the Caucasian population (http://allelefrequencies.net). Through IVS of PBL from an HLA-A*11:01 donor, we detected, isolated, and expanded T cells specifically recognizing three nonC-TL (Figure 4B). Two peptides mapped to 5’-UTR regions of the canonical genes HOXC13 (5’U-HOXC13) and ZKSCAN1 (5’U-ZKSCAN1), and one peptide mapped to a non-coding spliced variant of C5orf22C gene (nc-C5orf22C, Supplemental Figure 11). The ORF encoding these nonC-TL were considerably short, with up to 21, 46, or 49 amino acids respectively, as confirmed by the recognition of APC electroporated with RNA encoding the predicted ORF (Supplemental Figure 12). Additionally, the loss of recognition of TCL transduced with Cas9-sgRNA targeting the 5’UTR HOXC13 genomic locus unequivocally confirmed the specificity of the T cells for this antigen and the exact genomic location from which it is transcribed and translated (Supplemental Figure 13).

**Figure 4.**
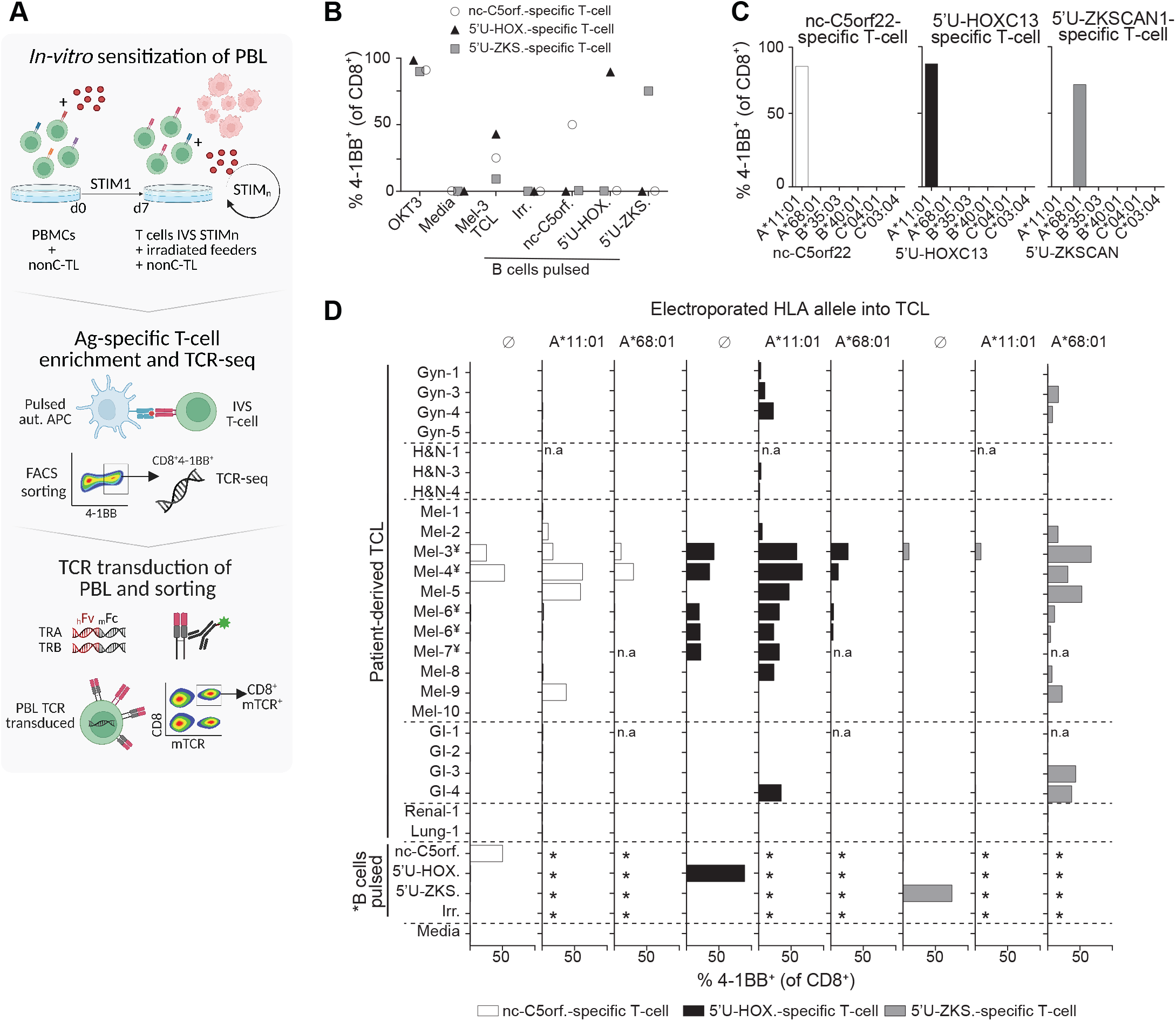
*In vitro* sensitization of donor PBL identified three immunogenic nonC-TL shared across patient-derived TCLs. (**A**) Donor PBL were in vitro sensitized (IVS) via three consecutive rounds of stimulation with 170 selected nonC-TL predicted to bind to HLA-A*11:01. Reactive T cells were enriched through FACS sorting based on CD8^+^ 4-1BB^+^ expression after 20 h co-culture with autologous B cells pulsed with the specific peptides and expanded for 14 days. The top 1 αβ pairs were cloned into a retroviral vector to trans-duce PBL and CD8^+^ mTCRB^+^ cells were FACS sorted to obtain a pure transduced population. Image created with BioRender. (**B**) Reactivity of antigen-specific T cells generated by IVS following FACS sorting enrichment. Frequency of 4-1BB^+^ on CD8^+^ cells after 20 h co-culture with B cells pulsed with the HPLC peptides specified is depicted. (**C**) Restriction element was evaluated by co-culturing enriched T-cell populations with COS-7 cells expressing the donor HLA alleles and pulsed with the corresponding peptides. (**D**) Expression and translation of the immunogenic nonC-TL in multiple patient-derived TCL indirectly evaluated through the detection of 4-1BB expression of nonC antigen-specific T cells co-cultured with TCL left untreated or electroporated with RNA encoding the specified HLA-I alleles. *B cells were not electroporated. ¥TCL naturally expressing HLA-A*11:01. n.a, non-assessed.

Although these three immunogenic nonC-TL were originally detected in Mel-3 TCL through immunopeptidomics, we investigated whether these antigens could be also expressed in other tumor cell lines. We exploited the high sensitivity of the nonC antigen-specific T cells identified to evaluate the expression and translation of these nonC antigens in a panel of 24 patient-derived TCL. T cells were cocultured with TCL artificially expressing the restriction element of interest (i.e., HLA-A*11:01 for 5’U-HOXC13 and nc-C5orf22, and HLA-A*68:01 for 5’U-ZKSCAN1, Figure 4C), in addition to the endogenous HLA alleles. Strikingly, we found that the three nonC-TL evaluated were frequently expressed and detected by nonC-TL-specific T cells in patient-derived TCL, as observed by 4-1BB upregulation when the relevant HLA was expressed (Figure 4D). Of note, 5’U-HOXC13 and 5’U-ZKSCAN1 were detected in melanoma, but also in other less immunogenic tumor types such as gastrointestinal cancers (GI) or gynecological malignancies. Furthermore, the recognition of several HLA-A*11:01^+^ TCL by nonC-TL-specific T cells without transfecting any additional HLA, showed that nc-C5orf22 and 5’U-HOXC13 can be naturally processed and presented on HLA-I (Figure 4D). Altogether, these results show that nonC-TL are shared across tumor types and can be naturally presented and recognized by T cells.

### TCRs targeting nonC antigens can display cancer-specific recognition

A potential concern that has not yet been addressed regarding the therapeutic targeting of nonC-TL is whether they are tumor specific. To investigate this, we analyzed the RNA expression of the canonical genes encoding for the three immunogenic nonC-TL in several solid tumors and matched healthy tissues from repository data (GEPIA). We compared their expression pattern to PMEL and MLANA as examples of melanoma-associated antigens, and MAGEA3 and MAGEC2 representing CGA (Figure 5A). Whereas C5orf22 and 5’U-ZKSCAN canonical genes displayed a variable but ubiquitous expression among tissues, the expression of HOXC13 canonical gene in healthy tissues appeared to be restricted to melanocytes, resembling the expression pattern of MLANA. This raised the possibility that the identified immunogenic nonC-TL might not be tumor specific. However, RNA transcript level cannot distinguish canonical from aberrant translation. In addition, contrary to the RNA-seq data analysis, the healthy immunopeptidome data that was used to select for nonC HLA-I ligands derived from ORFs absent in non-malignant cells suggested that the nonC translation of these peptides was tumor specific.

**Figure 5.**
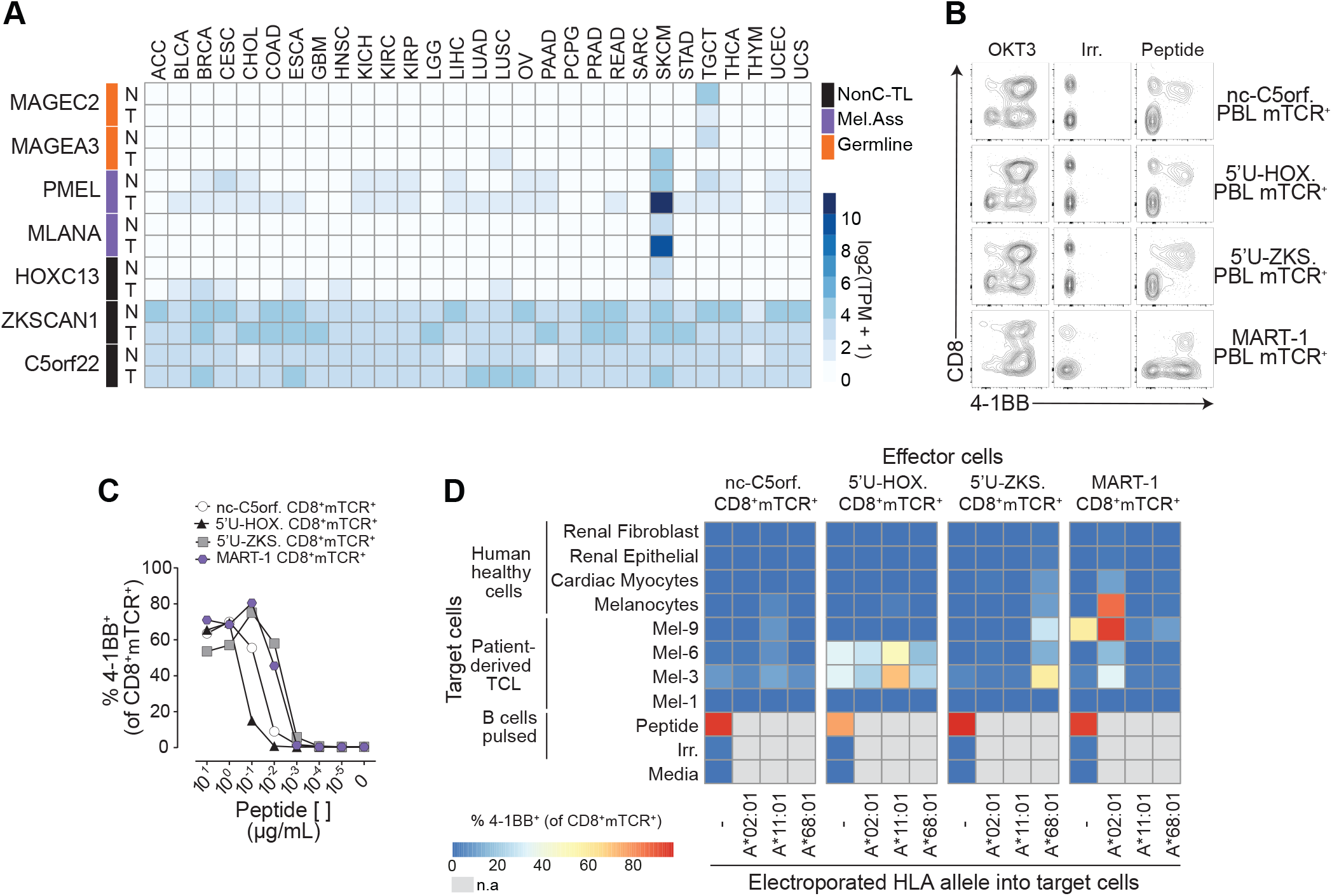
Evaluation of the tumor specificity of the three immunogenic nonC-TL. (**A**) RNA expression analysis in tumors (T) and matched healthy tissues (N) of the canonical genes encoding immunogenic nonC-TL compared to TAA and CGA. TCGA and GTEX data were obtained from GEPIA. (**B**) CD8 coreceptor activation-dependence of PBL transduced with antigen-specific TCRs. FACS plots show the expression of 4-1BB by CD8 (gated on CD3^+^mTCR^+^) after co-culture with peptide pulsed B cells. B cells pulsed with an irrelevant (Irrel.) peptide were used as negative control. (**C**) TCR-transduced T cells purified by FACS sorting (CD8^+^mTCR^+^) were co-cultured with B cells pulsed with serial dilutions of the corresponding peptide. SD mean is plotted.(**D**) Expression and translation of nonC-TL in healthy human cells and selected TCLs was indirectly evaluated by co-culturing control and electroporated target cells with RNA encoding the specified HLA alleles with sort purified TCR-transduced T cells. T-cell activation was assessed by measuring 4-1BB expression on CD8^+^mTCR^+^. n.a, non-assessed.

To gain further insights into the selective expression of these nonC-TL in tumor cells and their applicability as targets for cancer immunotherapy, we empirically evaluated the expression and translation of the selected immunogenic nonC-TL in several human healthy cell types by exploiting the ability of antigen-specific T cells to detect their cognate peptides with high sensitivity. To this end, we first sequenced the TCR locus of the nonC-TL-specific T cells identified by IVS and the most frequent TCR-α/β pairs of each of the populations were cloned into a retroviral vector and used to transduce PBL. Additionally, given that the canonical HOXC13 gene expression pattern resembled MLANA, we generated DMF5 TCR-transduced cells as an example of a TCR tested in clinical trials, which recognizes the melanomaassociated antigen MART-1_27-35_ (encoded by MLANA and restricted to HLA-A*0201) that is expressed both in melanoma cells and healthy melanocytes (8). Given that the TCRs recognizing nonC-TL were CD8-dependent, as opposed to the MART-1-specific TCR (Figure 5B), all PBL TCR transduced cells were sorted based on CD8 and mTCR expression (CD8^+^mTCR^+^). Peptide titration experiments to measure functional avidity of TCR transduced cells evidenced that nc-C5orf22 and 5’U-HOXC13-specific TCRs required higher concentrations of minimal peptide to become activated, while 5’U-ZKSCAN1 and MART-1 TCR transduced cells were more sensitive at detecting their cognate antigen (Figure 5C).

Next, the four antigen-specific CD8^+^ mTCR^+^ cells were co-cultured with human cells of different origin, including normal melanocytes, cardiac myocytes, renal epithelial, and fibroblast, as well as a few of the previously tested melanoma cell lines. As in the previous experiment used to evaluate the expression of nonC-TL antigens in patient-derived TCL (Figure 4D), expression of the specific HLA alleles restricting antigen recognition of the TCRs evaluated was exogenously enforced in the target cells and T-cell activation was evaluated by measuring 4-1BB upregulation in mTCR^+^ cells by flow cytometry following co-culture (Figure 5D and Supplemental Figure 14). As expected, MART-1 TCR transduced cells strongly recognized human melanocytes electroporated with the HLA-A*02:01, as well as three out of the four patient-derived melanoma cell lines tested. Surprisingly, cardiac myocytes were also recognized by MART-1 TCR transduced cells, albeit to a limited extent. 5’U-ZKSCAN1 TCR transduced cells displayed preferential recognition of melanoma cells compared to normal melanocytes and cardiac myocytes when target cells were electroporated with the relevant allele HLA-A*68:01. Importantly, 5’U-HOXC13 displayed recognition of two of the four melanoma cell lines included but did not recognize cardiac myocytes and barely recognized normal melanocytes when electroporated with the corresponding allele HLA-A*11:01. The lower sensitivity of the nonC C5orf22 TCR and/or limited expression of the antigen precluded us from reaching a conclusion regarding the tumor specific expression of this antigen. Overall, these results indicate that the 5’U-HOXC13 peptide is the nonC antigen with the highest tumor specificity followed by 5’U-ZKSCAN1. Importantly, both nonC antigens displayed a superior tumor specific profile compared to MART-1. Although we cannot rule out the possibility that these antigens could be expressed and translated in additional normal cell types, our findings suggest that the aberrant translation giving rise to the nonC peptides studied occurs preferentially in tumor cells rather than in normal cells. Overall, our results demonstrate that nonC-TL are a promising alternative source of tumor antigens to neoantigens, CGA and TAA not only because they can be naturally presented but also because they can be immunogenic and expressed across diverse tumor types but not, or at very low levels, in healthy cells.

## Discussion

Tumor antigens play an integral role driving protective or therapeutic antitumor immunity. Recent evidence suggests that peptides derived from nonC proteins can be systematically detected through proteogenomics and are specifically presented on HLA-I by tumor cells. However, their contribution to tumor immune surveillance and their immunogenicity has not been explored in detail.

The proteogenomics pipeline we used, Peptide-PRISM, is independent of RNA-seq and Ribo-seq but enables the detection of HLA-I ligands potentially originating from any region of the genome, including CDS, UTR, off-frame, ncRNA, intronic and intergenic regions (Figure 1A). Like previous immunopeptidomics studies (20–24,31,33,34,39), we observed that nonC HLA-I ligands were frequently detected across different cancer types (Figure 1B). Because we were interested in tumor-specific candidates we used healthy immunopeptidome data to exclude peptides derived from ORFs present in non-malignant cells. As a result, 61.5% of the nonC HLA-I-ligands detected were preferentially presented by tumors, referred to as nonC-TL (Figure 1C). Importantly, we studied the repertoire of presented tumor antigen candidates in 9 patient-derived TCL. Our work shows that nonC-TL (n=507) outnumber the HLA-I ligands derived from conventional tumor antigens such as mutations (n=33), CGA antigens (n=36) and melanoma-associated antigens (n=24) (Figure 2A). In line with previous reports (36–38), the number of HLA-I ligands derived from NSM was relatively low compared to the NSM identified by WES (Figure 2C). Notably, our data adds a considerable number of mutated HLA-I peptides to those identified through immunopeptidomics so far, underscoring the performance of Peptide-PRISM at detecting neoantigen candidates.

NonC proteins were frequently presented in patient-derived TCLs, being the main source of candidate tumor antigens, but pre-existing T-cell responses targeting nonC-TL were not detected in any of the patients studied (Figure 3F and Supplemental Figure 15). In contrast, nearly 65% of the antigens recognized by T cells were derived from mutations (n=13), 20% from CGA (n=4) and 15% from melanoma-associated antigens (n=3) (Table 1 and Supplemental Figure 15). Moreover, the isolated antigen-specific T cells recognized the autologous TCL (Supplemental Figure 7 and Figure 10), demonstrating that the peptides identified were bona fide tumor antigens. To our knowledge, the existence and frequency of naturally occurring T cells targeting nonC-TL compared to conventional tumor antigens has not previously been investigated in such detail. The majority of studies identifying nonC peptides presented on HLA-I did not investigate their immunogenicity in patients (20,22–24,31,33,40). A few explored their immunogenicity through *in vitro* sensitization of PBL from healthy donors which detects naïve rather than antigen-experienced T cells (21,41–43), or immunized mouse models (25). Only one proteogenomics report evaluated T-cell responses against nonC HLA-I ligands as well as other relevant tumor antigens derived from melanoma-associated antigens and CGA identified from patient-derived TCL and tumor samples(34). Although they reported some degree of reactivity to one of the 571 nonC HLA-I peptides evaluated, our results are in accordance with their findings since the percentage of reactive peptides clearly favored melanocyte differentiation antigens, supporting that nonC HLA-I ligands are not as immunogenic.

One limitation of our study lies in the nature of the proteogenomics approach used combined with the stringent and uniform 1% FDR threshold set to select the tumor antigen candidates. Although this pipeline potentially detects peptides originating from any region of the genome, the FDR calculation using a stratified mixture model could result in an underestimation of nonC ligands with an increased search space, for example intergenic regions. Consequently, some of the previously described nonC sources were not properly interrogated such as endogenous retroviral elements (ERE), or not considered in this study such as RNA editing or peptide splicing (44,45). Overall, our findings predict a limited contribution of nonC-TL to tumor immune surveillance.

Despite we could not detect recall T-cell responses to nonC-TL, we were able to isolate and expand T cells specifically targeting three nonC-TL through IVS of non-autologous HLA-A*11:01^+^ PBL. We found that two immunogenic nonC-TL derived from the aberrant translation of 5’UTR of HOXC13 and ZKSCAN1 genes were frequently detected in patient-derived melanoma TCL as well as other less immunogenic tumor types such as GI or Gyn. In addition, an immunogenic nonC-TL derived from a non-coding spliced variant of C5orf22 gene was also detected in several melanoma TCL. These results revealed that nonC-TL can be immunogenic and shared across tumor types, thus representing attractive targets for off-the-shelf vaccines or T-cell therapies.

The paradoxical lack of recognition of nonC-TL in cancer patients in light of the fact that at least a fraction of these were proven to be immunogenic could be explained through different mechanisms. For instance, we screened *ex vivo* expanded lymphocytes which frequently present a skewed oligoclonal TCR repertoire, which could lead to the depletion of some tumor-reactive clones (46). Although this could be the case, theoretically, this would negatively impact on the T-cell recognition of all antigen categories equally. Another possible explanation is that the level of expression of nonC-TL may be sufficient for presentation on tumor HLA-I, but inadequate for efficient cross-presentation *in vivo*, leading to defective priming of T cells. In line with this hypothesis, nonC peptides are thought to be less abundant and largely originated from disordered or unstable proteins with shorter half-lives compared to functionally annotated proteins (22,47). In addition, some nonC ORF are so small that they do not require processing (33). These characteristics can facilitate the accessibility of nonC peptides into the HLA-I antigen presentation pathway (47,48). Moreover, some studies have demonstrated that rapidly degraded proteins and minimal epitopes are unable to provoke cross-priming by APC as opposed to stable, full-length antigens (49). Altogether, this data suggests that the native characteristics of the aberrant translation events giving rise to nonC-TL (i.e. low abundance and instability) could account, at least in part, for the lack of recognition in cancer patients. An alternative explanation for our findings is that nonC-TL could also be presented by mTEC cells in the thymus, leading to partial central tolerance and, consequently, limiting the abundance of T cells targeting nonC-TL in periphery. This, combined with the low level of expression and/or poor priming could hamper the detection of antigen-experienced T cells targeting nonC-TL in cancer patients.

In our work, we addressed the tumor specific expression of nonC antigens, an essential aspect for the development of immunotherapeutic interventions. Previous studies have used healthy RNA-seq data to exclude nonC HLA-I ligands presented in healthy tissues. Alternatively, Ribo-seq could potentially be used to select tumor-specific nonC ORF. However, this technique is relatively new, the sensitivity is still limited and little data from healthy tissue is currently available. Instead, we leveraged a healthy immunopeptidome dataset to select nonC HLA-I ligands absent in non-malignant cells. Although immunopeptidomics is less sensitive, we believe it is more relevant, since it can detect peptides derived from both canonical and aberrant translation. To gain additional insights into the tumor specificity of the three immunogenic nonC identified, we indirectly evaluated their expression and translation in healthy human cells derived from several vital organs as well as melanocytes by using lymphocytes transduced with antigen specific TCRs. Our results revealed that the 5’U-HOXC13 peptide was the nonC antigen with the highest tumor-specificity, since TCR-transduced T cells targeting this antigen recognized several TCL, but barely melanocytes or any other healthy human cell tested. In fact, both 5’U-HOXC13 and 5’U-ZKSCAN1 nonC antigens displayed a far superior tumor specific profile compared to MART-1, as TCR DMF5 displayed strong recognition of melanocytes (>90%). Unexpectedly, DMF5 TCR transduced T cells recognized cardiac myocytes at a similar level as TCR transduced T cells targeting the nonC-TL 5’U-ZKSCAN1 (10-20%). Although DMF5 TCR has shown objective tumor responses in patients it also evidenced severe on-target toxicities in skin, eyes, and ears due to the high expression of MART-1 in melanocytes(8), but toxicities related to cardiac myocyte recognition has never been reported. These findings suggest that the low level of expression of these nonC-TL in healthy tissues that we observed (Figure 5D) are insufficient to trigger undesired off-tumor toxicities. Overall, our data supports that nonC-TL can display a tumor-specific profile that is compatible with their use as therapeutic targets. Nonetheless, these data must be interpreted with caution, since we cannot rule out the possibility that these immunogenic nonC-TL are expressed and translated in other healthy cells.

Overall, our results demonstrate that nonC-TL constitute an abundant source of candidate tumor antigens compared to peptides derived from mutations, CGA, or tissue differentiation antigens. NonC-TL are shared across tumor types and can be naturally presented by cancer cells and recognized by T cells. More importantly, they are not detected or at very low levels in healthy cells. Therapeutic interventions such as vaccines or TCR gene engineered T cells targeting nonC-TL could overcome the defective endogenous T cell responses observed in cancer patients by enhancing the *de novo* priming of T cells or administering large numbers of effector cells that can directly attack tumor cells. In fact, it is tempting to speculate that the lack or defective T-cell response to nonC tumor antigens in cancer patients could be advantageous, since these antigens and the HLA presenting them are less exposed to immune selective pressure and, consequently, tumors would less likely present antigen escape variants selected through immunoediting. Our findings predict a limited contribution of nonC-TL to cancer immunosurveillance and, at the same time, underscore nonC-TL as a promising source of antigens for the development for therapeutic interventions.

## Methods

### Patient characteristics

Patients Gyn-1, Gyn-2, Gyn-3, Gyn-4, H&N-1, H&N-3, Mel-1, Mel-2 and Mel-3 were chosen for this study on the basis of availability of autologous tumor cell line and matched lymphocytes. Patient characteristics are summarized in Supplemental Table 1.

### Establishment of patient-derived TCL

A small fragment (2-4 mm^3^) of tumor biopsies or surgically resected tumor was cultured in RPMI 1640 plus (Lonza) containing 10% FBS Hyclone (GE Healthcare), 100 U/mL penicillin (Lonza), 100 μg/mL streptomycin (Lonza) and 25 mM HEPES (Thermo Fisher Scientific) at 37°C in 5% CO2. The medium was replaced once every month until the TCL was established and then further expanded in T2 media containing RPMI 1640 plus (Lonza), 10%-20% FBS (Gibco), depending on the TCL, 100 U/mL penicillin (Lonza), 100 μg/mL streptomycin (Lonza) and 25 mM HEPES (Thermo Fisher Scientific) or cryopreserved until used. TCL were regularly tested for mycoplasma and were authenticated based on the identification of patient-specific somatic mutations and HLA molecules.

### TIL expansion

Small tumor fragments (2-4 mm^3^) were cultured in individual wells of a 24-well plate in T-cell media consisting of RPMI 1640 plus (Lonza) supplemented with 10% human AB serum (BST), 100 U/mL penicillin (Lonza), 100 μg/mL streptomycin (Lonza), 2 mM L-Glutamine (Lonza), 25 mM HEPES (Thermo Fisher Scientific) and 6e6 IU IL-2 (Proleukin) at 37°C and 5% CO2. Fresh media containing IL-2 was added on day 5 and media was changed, or TIL were split when confluent every other day thereafter. T cells were expanded independently for 15-30 days and cryopreserved until use. In some cases, T cells underwent a rapid expansion protocol (REP), as explained below.

### Rapid expansion protocol (REP)

T cells were expanded for 14 days using 30 ng/mL anti-CD3 (OKT3, Biolegend), 3e3 IU/mL of interleukin IL-2 (Proleukin) and irradiated allogeneic PBMC (50 Gy) pooled from three donors as feeder cells in T-cell medium RPMI 1640 plus (Lonza):AIM-V (Gibco) containing 5% human AB serum (BST), 100 U/mL penicillin (Lonza), 100 μg/mL streptomycin (Lonza), 2mM L-Glutamine (Lonza), 12.5 mM HEPES (Thermo Fisher Scientific). After day 6, half of the medium was replaced with fresh T-cell medium containing IL-2 every other day. Cells were split when confluent, harvested on day 14, and cryopreserved until use.

### PBMC isolation

Peripheral blood mononuclear cells (PBMC) were obtained using a Ficol density gradient (Lymphoprep, Stem cell) from pheresis or whole blood and cryopreserved for cell sorting, DNA extraction for WES and to expand B cells *ex vivo*.

### T cell sorting from PBL

PBLs were sorted based on the expression of surface markers previously described to enrich for tumorreactive T cells in peripheral blood such as PD-1 (50). Briefly, PBMCs were thawed and rested overnight without cytokines. Following CD8^+^ enrichment using CD8 microbeads (Miltenyi Biotec), the Fc receptor was blocked (Miltenyi Biotec) and cells were stained with the following antibodies for 30 minutes at 4°C: CD3PECy7 (BD, clone SK7, 0.5:50), CD8-APCH7 (BD, clone SK1, 1:50), PD1-PE (Biolegend, clone EH12.2H7, 0.75:50), CD38-APC (Biolegend, clone HIT2, 0.5:50) and HLA-DR BV605 (Biolegend, clone L243, 0.75:50). CD3^+^CD8^+^ cells expressing PD1hi alone or in combination with HLA-DR and CD38 were sorted in BD FACS AriaTM and expanded using a REP as previously specified.

### Generation of autologous APC

B cells were isolated from cryopreserved PBMCs by positive selection using CD19^+^ microbeads (Miltenyi Biotec) and expanded through CD40-CD40L stimulation by culturing cells for 4-5 days with irradiated NIH3T3 feeder cells constitutively expressing CD40L at 37°C in 5% CO2 in B cell medium. Iscove’s IMDM media (Gibco) containing 10% human AB serum (Biowest), 100U/mL Penicillin and 100 μg/mL streptomycin (Lonza), 2 mM L-Glutamine (Lonza), and supplemented with 200 U/ml IL-4 (Peprotech). Up to three rounds of stimulation and expansion were performed consecutively. B cells were cryopreserved from day 5 to 6 until use. When used after cryopreservation, B cells were thawed in B cell medium containing DNAse (Pulmozyme, Roche) 20 h before use in co-culture assays. Alternatively, CD4^+^ T cells were isolated from PBMCs by positive selection using CD4^+^ microbeads (Miltenyi Biotec) or FACS sorting and subsequently expanded through a REP.

### Peptides

The amino acid sequences of the identified tumor antigen HLA-I ligands were purchased from JPT Peptide Technologies (Berlin, Germany) as crude and used for screening, IVS, and MS validation with synthetic peptides. HPLC peptides were supplied by JPT Peptide Technologies (Berlin, Germany) and used in coculture experiments to confirm the reactivities. Selected endogenous HLA-I ligands were ordered from Thermo Fisher Scientific as crude (PePotec grade 3) with one stable isotope-labeled amino acid and used for PRM validation (See Supplemental data 3).

### Cloning, *in vitro* transcription of RNA, and electroporation

The HLA sequences of interest or predicted ORF of the immunogenic nonC-TL peptides were cloned into pcDNA3.1 using BamH1 and EcoR1 containing a Kozak motif upstream of the start codon. HLA-I sequences were obtained from IPD-IMGT/HLA and codon-optimized. The predicted ORF were constructed from the second nearest upstream in-frame start codon (ATG, CTG, or GTG) to the first in-frame stop codon downstream; the sequence was not codon optimized nor additional start codons were added. All the plasmids were synthesized by Genscript.

For in vitro transcription (IVT) of RNA the plasmids were linearized with Not-I followed by phenolchloroform extraction and precipitation with sodium acetate and ethanol. Next, 1 μg of DNA was used as a template to generate RNA by IVT using HiScribe# T7 ARCA mRNA Kit with tailing (New England) following manufacturer’s instructions. RNA was precipitated using LiCl2, resuspended at 1 μg/μL in molecular grade H2O, and stored at -80º until use.

From 0.5-1e6 TCL, healthy human cells, and B cells were harvested and resuspended in 100 μl of Opti-MEM media (Gibco) and transferred into a sterile 0.2 cm cuvette (VWR electroporation cuvettes). From 4 to 8 ug of RNA encoding for the sequence of interest were added for electroporation. Cells were electroporated at 150 V, 20 ms, and 1 pulse using an ECM 830 BTX-Electroporator. After electroporation, cells were resuspended in pre-warmed specific media containing DNAse (Pulmozyme, Roche). After 20 h cells at 37ºC and 5% CO2, cells were harvested, washed with PBS, and used in co-culture assays. A GFP RNA electroporation control was included for each cell line and assessed by Flow cytometry as a transfection control.

### Co-culture assays: IFN-γ enzyme-linked immunospot (ELISPOT) assays and detection of activation marker 4-1BB using flow cytometry

T cells were thawed into T-cell medium supplemented with 3,000 IU IL-2 (Proleukin) and DNAse (Pulmozyme, Roche) three to four days before coincubation with target cells. All co-cultures were performed in the absence of exogenously added cytokines. Cells were stained with CD3-APCH7 (BD, clone SK7, 0.3:40), CD8-PECy7 (BD, clone RPA-T8, 0.1:40), CD4-PE (BD, RPA-T4, 0.3:40) and CD137-APC (BD, clone 4B4-1, 0.5:40), and in some cases mTRB-FITC (eBiosciences, clone H57-597, 0.2:40) for 30 minutes at 4°C, washed with staining buffer containing PI (1:2000) and acquired in BD FACSLyric™, BD FACSCanto™ or BD FACSLyric™. In parallel, IFN-y secretion was detected using IFN-γ capture and detection antibodies (MABtech technologies) assessed by ELISPOT assay following manufacturer instructions. ELISPOT plates were analyzed and counted in ELISPOT reader. For all the assays, plate-bound OKT3 (1 μg/mL; Biolegend) was used as a positive control. Media, and/or autologous APC pulsed with irrelevant peptides were used as negative controls.

For the detection of recall T-cell responses, from 2e4 to 5e4 *ex vivo* expanded TIL, sorted PBL or enriched populations of tumor-reactive lymphocytes were co-cultured with 1e5 to 2e5 peptide-pulsed autologous APC (either B cells, or CD4+ T cells). T-cell reactivities were considered positive if the number of IFN-γ spots were greater than double the amount of the irrelevant control condition and greater than 40 spots. Additionally, reactivities had to be observed in at least two independent experiments. Crude peptides preparations were used for screening, and the reactivities were further confirmed with HPLC grade peptides. Experiments were performed at least twice.

### Enrichment of tumor-reactive and antigen-specific T cells

Either expanded TIL, sorted PBL or IVS T cells were co-cultured with tumor cells or peptide-pulsed autologous APC for 20 h. CD3^+^CD8^+^ cells expressing 4-1BB were sorted in BD FACS AriaTM or BD Influx™ and expanded using a REP as previously specified. The same antibodies and dilutions used for co-cultures described above were scaled up for staining 4-1BB+ T cells.

### HLA restriction element determination

COS-7 cells were transfected with plasmids encoding the individual HLA molecules using Lipofectamine 2000 (Life Technologies). After resting overnight, cells were harvested and pulsed with the corresponding peptides for 2 h, washed, and used as targets in co-culture assays.

### *In vitro* sensitization of PBL

HLA-A*11:01 donor PBMCs were stimulated with 5 independent peptide pools (PP) each containing up to 35 nonC-TL selected by the prediction score to bind to HLA-A*11:01 according to NetMHCpan4.0. Cells underwent three consecutive rounds of stimulation every 7 days with 0.25 μg/mL per peptide and a combination of IL-21, IL-7 and IL-2. More specifically, at day 0, 5e6 donor PBMC were cultured in 24-well plates with OpTmizer™ media (Gibco) containing IL-21 (Peprotech 25 ng/mL) and the corresponding PP at 0.25 ug/ml per peptide. On day 6, IL-2 (Proleukin 18 IU/mL) and IL-7 (Peprotech 10 ng/mL) were added. For STIM2 (day 7) and STIM3 (day 14), T cells from the previous STIM were harvested, counted, and restimulated with autologous irradiated PBMC (50 Gy) pulsed with the corresponding PP at 1:10 ratio. Thereafter, fresh OpTmizer™ media (Gibco) containing IL-2 (18 IU/mL) and IL-7 (10 ng/mL) was replaced when medium looked acidified, or cells required splitting.

*De novo* T-cell responses were evaluated after three stims by co-culturing IVS T cells with autologous B cells pulsed with the corresponding PP and analyzing 4-1BB upregulation by flow cytometry as described above.

T cells recognizing the corresponding PP were sorted based on 4-1BB expression and expanded for 14 days in a REP (Enrichment of antigen-specific T cells). To identify the specific peptide recognized within the PP, sorted populations were co-cultured with B cells pulsed with individual peptides. The recognition was confirmed using HPLC purified peptides.

### TCR sequencing and PBL transduction

The TCR locus was sequenced by multiplex single-cell RNA sequencing of enriched antigen-specific T-cell populations. The samples were multiplexed using TotalSeq™ barcodes. Sequencing was done on an Illumina NS6000 with an S1 flowcell and v1 chemistry. Mapping, quantification, and clonotype definitions were done using cell ranger multi software (version 6.1.1 using the reference vdj_GRCh38_alts_ensembl-5.0.0). Demultiplexing and subsequent analysis was using the packages Seurat (version 4.0.3) and scRepertoire (version 1.3.5); Seurat::HTODemux was run using default parameters to obtain singlets.

TRA V-J-encoding sequences and TRB V-D-J-encoding sequences were combined to sequences encoding the mouse constant TRA and TRB chains (51), respectively. Mouse constant regions were modified, as previously described (52,53). The full-length TRB and TRA chains were cloned separated by a furin SGSG P2A linker into pMSGV1 retroviral vector (GenScript). Transient retroviral supernatants were generated by transfecting the vector encoding the TCR of interest (MSGV1) and envelope (RD114) into 293GP cells using Lipofectamine 2000 (Life Technologies). PBLs were activated in T cell medium supplemented with 50 ng/mL anti-CD3 and 300 IU/mL IL-2 for 3 days before retroviral transduction. Retroviral supernatants were harvested at 24 and 48 hours, centrifuged to discard cell debris, and diluted 1:1 with medium and used to transduce the activated lymphocytes using the spinoculation method, as previously described.(15)

### Normal human cell lines

Normal human cell lines were purchased from Promocell, thawed and cultured following manufacturer’s instructions in the recommended media without antibiotics. Cells were split when confluent with Dettaching kit (Promocell), cultured at the recommended concentration and expanded no more than 4 passages until use. HCM-c (Cat: C-128810) were cultured in myocyte growth medium (Cat nº: C-39275). HREpC-c (CatC-12665) were cultured in Renal Epithelial Cell GM media (Cat nº: C-39606). HSAEpC-c (Cat: C-12642) were cultured in Small Airway Epithelial cell GM (Cat nº: C-39175). NHEM.f-c (Cat: C-12400) were cultured in Melanocyte growth medium (Cat nº: C-39415).

### Whole exome sequencing

To identify the tumor-specific NSM, genomic DNA was purified from a cell pellet of patient-derived TCL and matched PBMC. WES libraries were generated by exome capture of approximately 20,000 coding genes using SureSelect human All exon V6 kit (Agilent Technologies) and paired-end sequencing was performed on a HiSeq sequencer (Illumina) at Macrogen. The average sequencing depth ranged from 100-150 for each of the individual libraries generated. Alignments of WES to the reference human genome build hg19 were performed using novoalign MPI from novocraft. Duplicates were marked using Picard’s MarkDuplicates tool. Insertion and deletion (indel) realignment and base recalibration were performed according to GATK best-practices. Samtools was used to create tumor and normal pileup files. Four independent mutation callers (Varscan, SomaticSniper, Mutect and Strelka) were used to call somatic NSM. The genomic coordinates from VCF files containing tumor-specific mutations were converted from hg19 to hg38 assemblies.

### GTEX and TCGA RNA analyses

TCGA and GTEX data from paired tumor and healthy data was obtained and analyzed using GEPIA (02/05/2022) and plotted using R. RNA levels are expressed as Log2 (TPM+1), the density of color in each block represents the median expression value of a gene in a given tissue, normalized by the maximum median expression value across all blocks. Abbreviation of tumor types: ACC;Adrenocortical carcinoma, BLCA;Bladder Urothelial Carcinoma, BRCA;Breast invasive carcinoma, CESC;Cervical squamous cell carcinoma and endocervical adenocarcinoma, CHOL;Cholangio carcinoma, COAD;Colon adenocarcinoma, DLBC;Lymphoid Neoplasm Diffuse Large B-cell Lymphoma, ESCA;Esophageal carcinoma, GBM;Glioblastoma multiforme, HNSC;Head and Neck squamous cell carcinoma, KICH;Kidney Chromophobe, KIRC;Kidney renal clear cell carcinoma, KIRP;Kidney renal papillary cell carcinoma, LAML;Acute Myeloid Leukemia, LGG;Brain Lower Grade Glioma, LIHC;Liver hepatocellular carcinoma, LUAD;Lung adenocarcinoma, LUSC;Lung squamous cell carcinoma, MESO;Mesothelioma, OV;Ovarian serous cystadenocarcinoma, PAAD;Pancreatic adenocarcinoma, PCPG;Pheochromocytoma and Paraganglioma, PRAD;Prostate adenocarcinoma, READ;Rectum adenocarcinoma, SARC;Sarcoma, SKCM;Skin Cutaneous Melanoma, STAD;Stomach adenocarcinoma, TGCT;Testicular Germ Cell Tumors, THCA;Thyroid carcinoma, THYM;Thymoma, UCEC;Uterine Corpus Endometrial Carcinoma, UCS;Uterine Carcinosarcoma,UVM;Uveal Melanoma.

### Purification of HLA-I peptides

Purified anti-HLA-I clone W6/32 (ATCC® HB95) antibodies were cross-linked to protein-A Sepharose 4B conjugate beads (Invitrogen) with dimethyl pimelimidate dihydrochloride (Sigma-Aldrich) in 0.2 M Sodium Borate buffer pH 9 (Applichem). From 5e7 to 3e8 tumor cells (See Supplemental Table 1) were snap-frozen, thawed, and lysed with PBS containing 0.6% CHAPS (Applichem) and Protease inhibitor Cocktail Complete (Roche). The cell lysates were sonicated (Misonix 3000) and cleared by centrifugation for 1 h at max speed to obtain the soluble fraction containing the pHLA complexes. The HLA-I affinity chromatography was performed using a 96-well single-use micro-plate with 3 μm glass fiber and 10 μm polypropylene membranes (Agilent). Sep-Pak tC18 100 mg Sorbent 96-well plates (Waters) were used for peptide purification and concentration as previously described (54). Peptides were eluted with 500 μl of 32,5% ACN in 0.1% TFA, lyophilized, and further cleaned and desalted with TopTips (PolyLC Inc.)

### LC-MS/MS acquisition

Acclaim Pep-Map nanoViper, C18 (Thermo Scientific) at a flow rate of 15 μl/min using a Thermo Scientific Dionex Ultimate 3000 chromatographic system (Thermo Scientific). Peptides were separated using a C18 analytical column of 75 μm × 250 mm, 1.8 μm, 100Å (Waters) or 25 μm × 250 mm, 1.8 μm, 100Å (Waters). Orbitrap Fusion Lumos™ Tribrid (Thermo Scientific) mass spectrometer was operated in data-dependent acquisition (DDA) mode. Survey MS scans were acquired in the orbitrap with the resolution (defined at 200 m/z) set to 120,000. The top speed (most intense) ions per scan were fragmented in the linear ion trap (CID) and detected in the Orbitrap with the resolution set to 30,000. Quadrupole isolation was employed to selectively isolate peptides of 400-600 m/z. Included charged states were 2 and 3. Target ions already selected for MS/MS were dynamically excluded for 10 s.

### Mass spectrometry data analysis of HLA-I peptides with Peptide-PRISM

Peptide-PRISM was used as previously described (20) without including random substitutions nor proteasome-spliced peptides. Briefly, for each identified fragment ion mass spectrum the Top 10 candidates were first identified by *de novo* sequencing with PEAKS X and later aligned to a database containing a 3-frame translated transcriptome (Ensembl90) and 6-frame translated genome (hg38). Additionally, vcf files from somatic mutation calling were used to interrogate NSM in a personalized fashion. All identified string matches were categorized into CDS (in-frame with annotated protein), 5’-UTR (contained in annotated mRNA, overlapping with 5’-UTR), Off-frame (off-frame contained in the coding sequence), 3’-UTR (all others that are contained in an mRNA), ncRNA (contained in annotated ncRNA), Intronic (intersecting any annotated intron) or Intergenic. Then, for each fragment ion mass spectrum, the category with the highest priority (CDS>5’UTR>Off-frame>3’UTR>ncRNA>Intronic>Intergenic) was identified, and all other hits among the 10 *de novo* candidates were discarded. The FDR was calculated for each category in a stratified mixture model considering the peptide length and database size. The same pipeline was applied to immunopeptidomics data obtained from HLA ligand atlas (55) including various tissues and HLA alleles. The predicted ORF from the nonC HLA-I ligands identified in the healthy immunopeptidome were retrieved and used to filter out the nonC HLA-I ligands from our tumor samples derived from the same ORF. All identified peptides were filtered to FDR 0.01. In addition, peptides with a *de novo* score (ALC) smaller than 30 and the sequences that could not be unequivocally assigned to a single category (Top location count=1) were filtered out (Supplemental Data 2). For peptides derived from NSM, the FDR was set at 0.02 (Supplemental Data 3).

### HLA-I typing and prediction of binding to patient-specific HLA molecules

HLA typing was determined from the WES data using the PHLAT algorithm (Supplemental Data 1). Eluted ligand likelihood (ELL) percentile rank scores for binding to the patient’s HLA molecules were obtained for all unique peptides ≥8 Aa eluted from TCL using NetMHCpan 4.0. The threshold for binding was set to <2%-tile rank.

### Validation of HLA-I peptides with synthetic peptides

Spectrum validation of the experimentally eluted HLA-I ligand tumor antigen candidates was performed by computing the similarity of the spectra acquired in the sample with the corresponding non-labeled synthetic peptide from the library. Briefly, synthetic crude peptides obtained from JPT were acquired in a pool using LC-MS/MS in conditions similar to those previously used to analyze samples to generate a spectral library. Peptide sequences were identified by database search with PEAKS-X Pro using a database containing Swiss-Prot as well as all the tumor antigen candidates interrogated. The search was exported as “for third party” format and imported into Skyline software to generate the library. The experimentally acquired HLA-I tumor samples were uploaded into Skyline and the similarity of the fragments (b/y ions) from the library (synthetic) vs. endogenous (sample) were analyzed considering library dot product (dotp) values, which range from 0 to 1 and, being dotp=1 the closest match. (See Supplemental Data 3).

### Validation of HLA-I peptides with isotope-labeled peptides

For each selected peptide, a synthetic isotope-labeled peptide at one chosen amino acid was spiked into the samples and used as an internal standard for Parallel Reaction Monitoring (PRM) detection. The amount of internal standard peptide to be spiked in each sample was evaluated using dilution curves and the final concentration was chosen based on a good chromatographic signal and no trace detectable of potential unlabeled traces from the synthetic internal standard. For Mel-3 TCL 30% of the sample (total of 5e7 cells) and for Mel-1 50% of the sample (total 1e8) was analyzed by PRM using different MS machines. Orbitrap Eclipse (Thermo Fisher Scientific) coupled to an EASY-nanoLC 1000 UPLC system (Thermo Fisher Scientific) with a 50 cm C18 chromatographic column. A PRM method was used for data acquisition with a quadrupole isolation window set to 1.4 *m/z* and MS2 scans over a mass range of *m/z* 250-1800, with detection in the Orbitrap mass analyzer at a 240 K resolution. MS2 fragmentation was performed using HCD fragmentation at a normalized collision energy of 30%, the AGC was set at 100,000, and the maximum injection time at 502 ms. All data were acquired with XCalibur software.

For data analysis fragment ion chromatographic traces corresponding to the targeted precursor peptides were evaluated with Skyline software v.21.2. Verification of the endogenous peptides was based on: i) the number of detected traces, ii) co-elution of endogenous traces, iii) co-elution of endogenous and internal standard peptides, iv) correlation of the fragment ions relative intensities between endogenous and internal standard peptides and, v) expected retention time.

### CRISPR/Cas9 Knock out

Single guides RNA (sgRNA) targeting the predicted ORF for 5’U-HOXC13 were designed with CRISPOR tool (http://crispor.tefor.net/). sgRNA specifically binding the genomic peptide location or upstream with the highest predicted KO efficiency were selected. Next, sgRNA were cloned into lenti-Cas9-v2 (Addgene, #52961) with BsmBI.

Lentiviral supernatants were generated by co-transfecting HEK293 cells with lenti-Cas9-v2 encoding the sgRNA of interest, psPAX2 and pMD2G plasmids with PEI (Sigma) and non-supplemented DMEM media (Gibco). Media was replaced after 12h with Opti-MEM (Gibco) containing 2% FBS (Gibco). Supernatants were collected at 42-48 h after transfection and filtered through a 0.45 um low-protein binding filter (Millipore). Patient-derived TCL were infected with lentiviral supernatants containing polybrene (Sigma) final concentration of 8 ug/mL followed by spinoculation 900 g, 32ºC, 50 min. Five days post-infection, T2 media (RPMI containing 10% FBS supplemented with Pen/Strep and L-Glut) was replaced with T2 media containing puromycin 1 μg/mL to select the cells efficiently transduced. Thereafter, media was replaced with fresh media containing puromycin when acidified or cells split when confluent. To evaluate the KO efficiency, puromycin-resistant cells were used as targets in co-culture assays with 5’U-HOXC13 specific T cells.

### Statistics

Experiments were performed without duplicates, unless otherwise specified, data were reported as mean +-SEM. All experiments were repeated at least twice. For HLA-I ligand identification with Peptide-PRISM, the FDR was calculated for each category in a stratified mixture model as previously described (20)considering the peptide length and database size.

### Study approval

Samples were obtained through a study approved by the Vall d’Hebron Hospital ethical committee (PR(AG)482/2017). All patients provided a written, informed consent.

## Data Availability

The mass spectrometry proteomics data have been deposited to the ProteomeXchange Consortium via the PRIDE [1] partner repository with the dataset identifier PXD036856. The source data underlying Figure 5A was downloaded from GEPIA (02/05/2022), data for Supplemental Figure 5 was downloaded from PRIDE (identifiers PXD022150 and PXD004894). All other data are available from the corresponding author on reasonable request.

## Author contribution

A.G. conceived and designed the project and interpreted the results and wrote the manuscript.

M.L.R. designed, performed the experiments, and interpreted the analyses and wrote the manuscript.

F.E. and A.S. conceptualized and implemented the software for MS/MS data processing and FDR calculations using Peptide-PRISM.

M.L.R. and A.Y.E. conducted the immunopeptidomics MS experiments.

M.L.R. generated patient reagents including TCL, TIL and sorted PBL. M.L.R., A.G.G. and J.P. conducted the TIL experiments.

F.C. assisted in MS experimental design, data analyses and visualization.

J.J.G. performed the NGS analyses.

M.G. provided support with MS immunopeptidomics

JM.L., M.O.O, I.M., A.V., JM.P., X.M.G, I.B., E.M., E.G., provided valuable patient reagents used in this study.

## Acknowledgements

We thank the patients for their participation in this study, Steven A. Rosenberg for providing valuable reagents and support for NGS studies, R. Pujol for helpful scientific discussion, J.Gonzalez for bioinformatics support, CRG/UPF Flow Cytometry Unit for assistance with cell sorting, and CRG/UPF and IRB Proteomics Units for technical support. A.G. and this work were funded by the Comprehensive Program of Cancer Immunotherapy & Immunology II (CAIMI-II) supported by the BBVA Foundation (53/2021), Institute Carlos III (MS15/00058 and PI17/01085), AECC (IDEAS197PORT), and La Fundació La Marató de TV3 (201919-30). We thank CERCA Programme / Generalitat de Catalunya for institutional support. M.L.R. was supported by the Agència de Gestió d’Ajuts Universitaris i de Recerca (AGAUR) (2018FI_B 00946). A.G.G was supported by Generalitat PERIS award (SLT017/20/000131). A.Y.E. was supported by the Agència de Gestió d’Ajuts Universitaris i de Recerca (AGAUR) (2021 FI_B 00365). J.P. was supported by the Beatriu de Pinós programme (BP 2018), cofounded by the Agency for Management of University and Research Grants (AGAUR) and European Union’s Horizon 2020.

## Supplemental Figures and tables

Supplemental Figure 1. Validation of HLA-I ligand tumor antigen candidates with synthetic peptides. Supplemental Figure 2. Validation of selected HLA-I ligand tumor antigen candidates identified in Mel-1 and Mel-3 with isotope-labeled peptides.

Supplemental Figure 3. Frequency of nonC peptides detected in HLA-I.

Supplemental Figure 4. HLA-I allele binding motif of all patients included in the study.

Supplemental Figure 5. A fraction of nonC-TL can be detected in publicly available immunopeptidomics datasets from human tumor biopsies.

Supplemental Figure 6. Evaluation of pre-existing T-cell responses to candidate tumor antigens in cancer patients.

Supplemental Figure 7. Enrichment of antigen-specific T cells from cancer patients by flow cytometry-based sorting.

Supplemental Figure 8. Deconvolution of peptide pools containing recognized CGA or melanoma-associated antigens.

Supplemental Figure 9. Functional avidity and specificity of neoantigen-specific T cells isolated from cancer patients.

Supplemental Figure 10. HLA restriction element and recognition of autologous tumor cell lines by antigen-specific T cells isolated from cancer patients.

Supplemental Figure 11. Genomic location of the three immunogenic nonC-TL identified through IVS. Supplemental Figure 12. Recognition of the predicted ORFs encoding the immunogenic nonC-TL by TCR transduced cells.

Supplemental Figure 13. Loss of recognition of TCL transfected with Cas9-sgRNA targeting the predicted 5’U-HOXC13 ORF.

Supplemental Figure 14. Assessment of expression and translation of the immunogenic nonC-TL in human healthy cells.

Supplemental Figure 15. NonC-TL are the main source of candidate tumor antigens identified by proteogenomics but tumor-reactive T cells preferentially recognize neoantigens derived from NSM. Supplemental Table 1. Patient characteristics and relevant experimental information.

## Supplemental data

Supplemental Data 1. HLA-I typing of tumor cell lines.

Supplemental Data 2. MS information HLA-I ligands FDR≤0.01 and ALC≥30. Supplemental Data 3. HLA-I ligands derived from NSM.

