## Supplemental Figures and Table for "Immunogenicity of non-canonical HLA-I tumor ligands identified through proteogenomics"

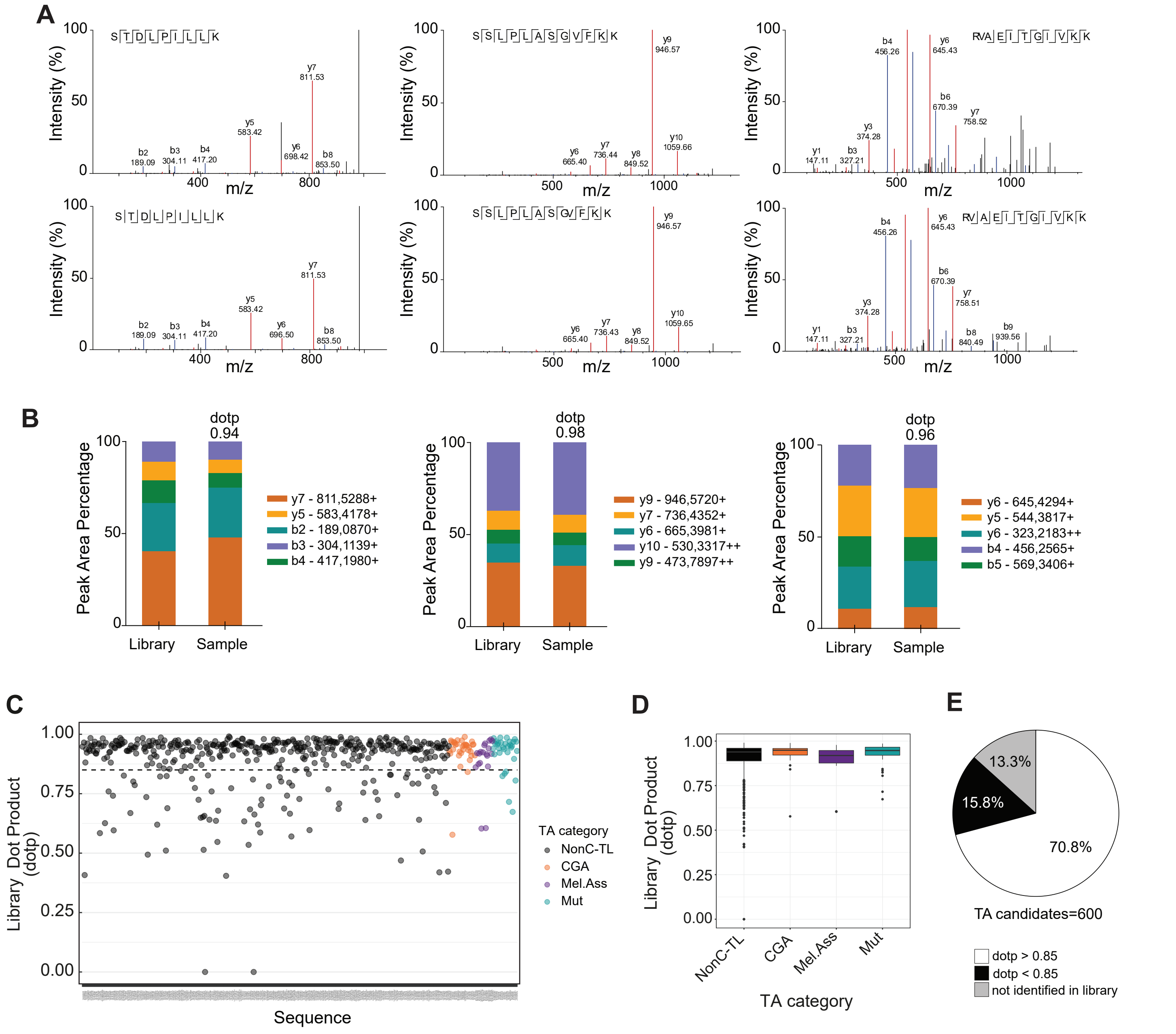


**Supplemental Figure 1. Validation of HLA-I ligand tumor antigen candidates with synthetic peptides.**
Spectrum validation of the experimentally eluted HLA-I ligand tumor antigen (TA) candidates was performed by computing the similarity of the spectra acquired in the sample with that of the corresponding non-labeled synthetic peptide from the library. A spectral library was generated with synthetic peptides acquired in a pool using comparable LC-MS/MS shotgun conditions. The similarity of the MS/MS fragmentation pattern from the library compared to the sample was evaluated using Skyline Software considering library dot product (dotp) values of the integrated peak areas for the observed fragment ions. (**A**) Representative examples of MS/MS fragmentation pattern of three nonC-TL eluted from Mel-3 TCL HLA-I (experimental identification, top) and their corresponding synthetic peptide (library, bottom), PEAKS XPro visualization. (**B**) b/y ions generated from the library (synthetic) compared to fragments acquired in the sample (experimental identification) with Skyline software. The similarity is measured according to dotp, which ranges from 0 to 1, being dotp=1 the closest match. Example of the same peptides represented in A. (**C**) Representation of the similarity between tumor antigen (TA) candidates identified in the experimental sample and their corresponding synthetic peptides. The analysis of the library dot product (dotp) (y-axis) is shown for each sequence identified sequence (x-axis). The threshold to consider a peptide validated was set at 0.85 dotp (dashed line). (**D**) Box and whisker plot for the dotp values for all the TA candidates identified in the library for each antigen category. (**E**) Percentage of candidates validated according to library dot product (dotp ≥0.85), not validated (dotp <0.85) or not interrogated (synthetic not detected, and thus not included in library).

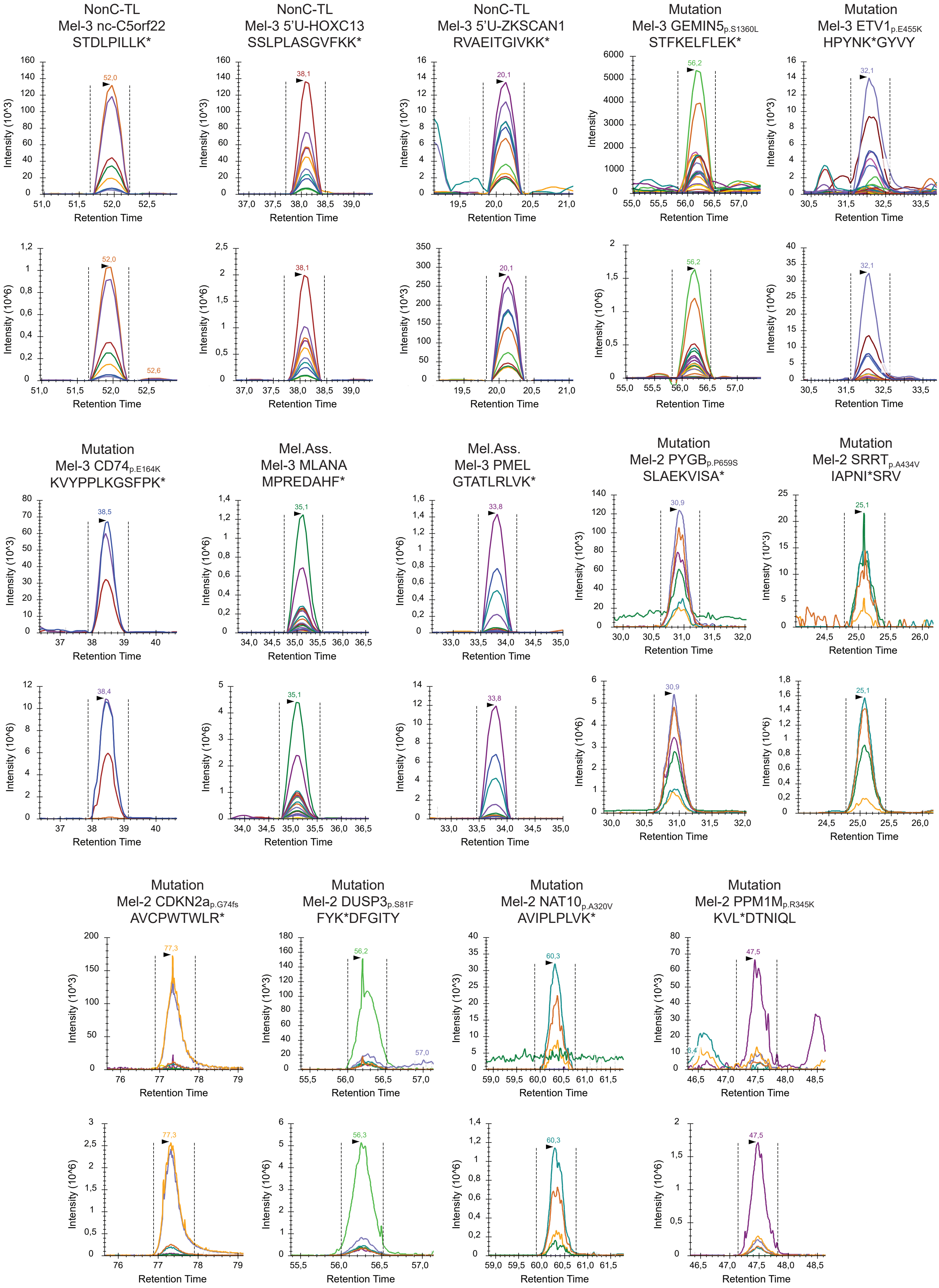
NonC-TL
Mel-3 nc-C5orf22
STDLPILLK*
NonC-TL
Mel-3 5'U-HOXC13
SSLPLASGVFKK*
NonC-TL
Mel-3 5'U-ZKSCAN1
RVAEITGIVKK*
Mutation
Mel-3 GEMIN5p.S1360L
STFKELFLEK*
Mutation
Mel-3 ETV1p.E455K
HPYNK*GYVY
Mutation
Mel-3 CD74p.E164K
KVYPPLKGSFPK*
Mel.Ass.
Mel-3 MLANA
MPREDAHF*
Mel.Ass.
Mel-3 PMEL
GTATLRLVK*
Mutation
Mel-2 PYGBp.P659S
SLAEKVISA*
Mutation
Mel-2 SRRTp.A434V
IAPNI*SRV
Mutation
Mel-2 CDKN2ap.G74fs
AVCPWTWLR*
Mutation
Mel-2 DUSP3p.S81F
FYK*DFGITY
Mutation
Mel-2 NAT10p.A320V
AVIPLPLVK*
Mutation
Mel-2 PPM1Mp.R345K
KVL*DTNIQL
Intensity (10^3)
Intensity (10^6)
Intensity
Retention Time

**Supplemental Figure 2. Validation of selected HLA-I ligand tumor antigen candidates identified in Mel-1 and Mel-3 with isotope-labeled peptides.**

Isotope-labeled peptide versions for each selected HLA-I ligand sequence were spiked into the corresponding HLA-I peptide sample and both the heavy and the light peptides were monitored through parallel reaction monitoring. Representative co-elution profiles of transitions of endogenous (top) and isotope-labeled (bottom) peptides using Skyline software are displayed for the HLA-I ligand tumor antigen candidates specified. Tumor antigen category, patient ID, and the amino acid sequence validated are specified for each peptide. Isotope-labeled amino acids are marked with *.

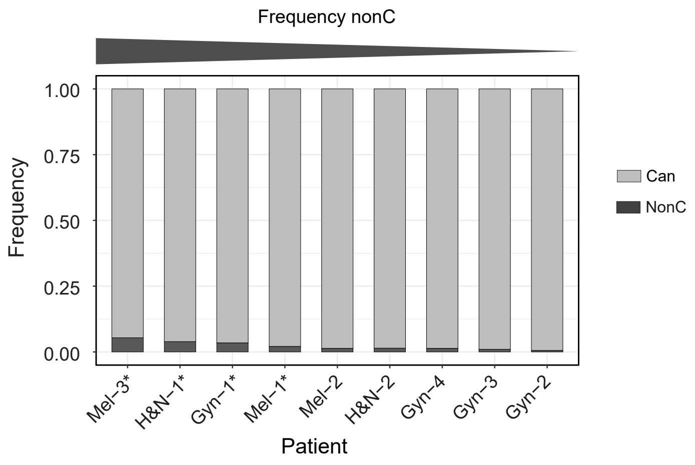


**Supplemental Figure 3. Frequency of nonC peptides detected in HLA-I.**
Percentage of peptides derived from canonical or non-canonical proteins for each patient ordered from higher to lower frequency of nonC HLA-I ligands. Patients harboring HLA-A*11:01 or HLA-A*03:01 are marked with an *.

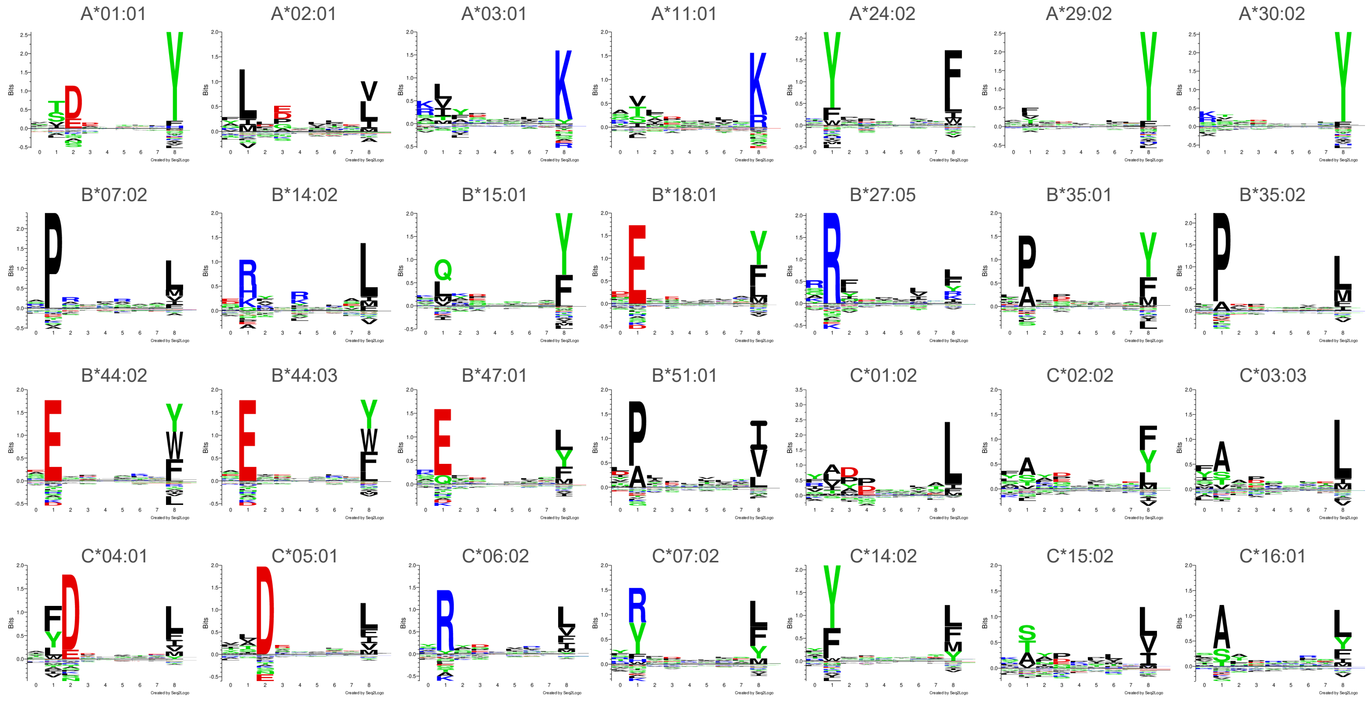

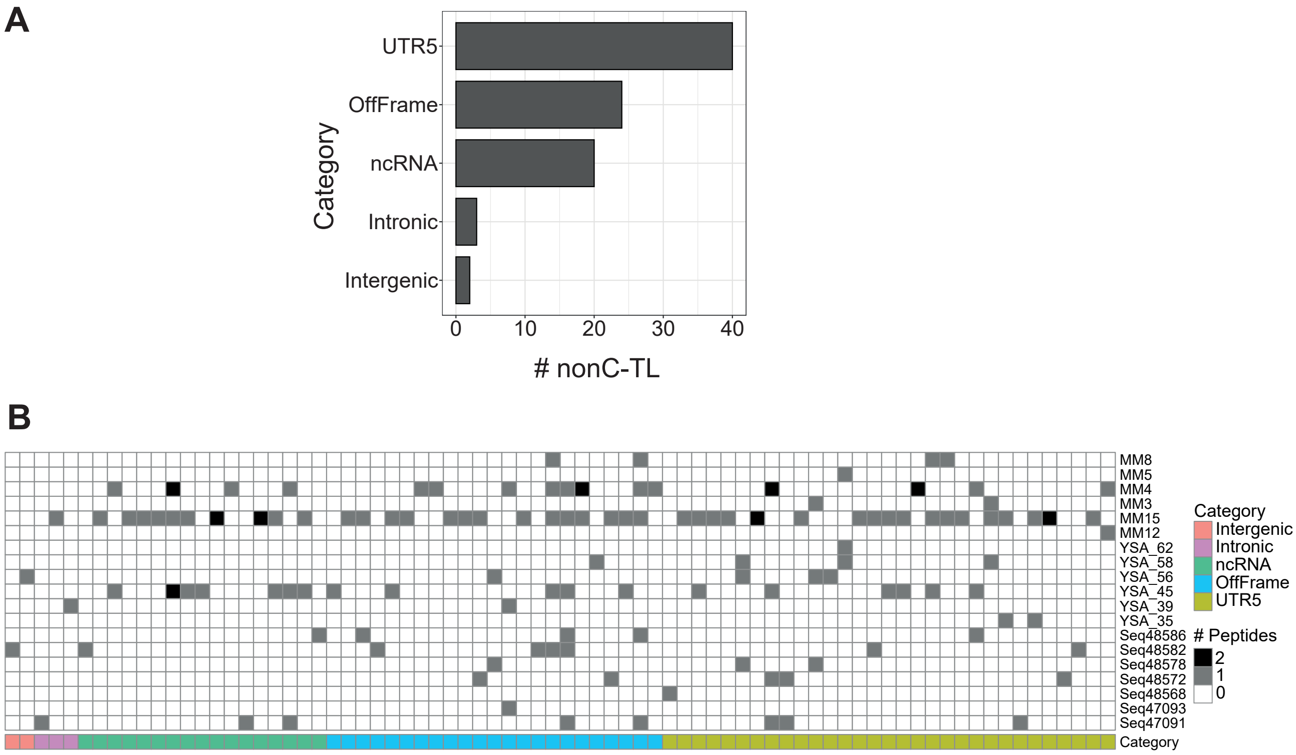


**Supplemental Figure 5. A fraction of nonC-TL can be detected in publicly available immunopeptidomics datasets from human tumor biopsies.**
HLA-I immunopeptidomics raw data of melanoma samples from Bassani-Sternberg et al., 2016 (36); Kalaora et al., 2021 (35), was downloaded from PRIDE and search against a database containing the reference proteome plus our nonC-TL candidates (n=507) using PEAKS Studio Pro. The analysis shows the nonC-TL sequences identified at 1% FDR. **a** Unique nonC-TL sequences identified by category. **b** NonC-TL identified in each sample. The tumor sample ID is shown on the right, and the nonC category and the number of times each peptide was detected is depicted in different colors.

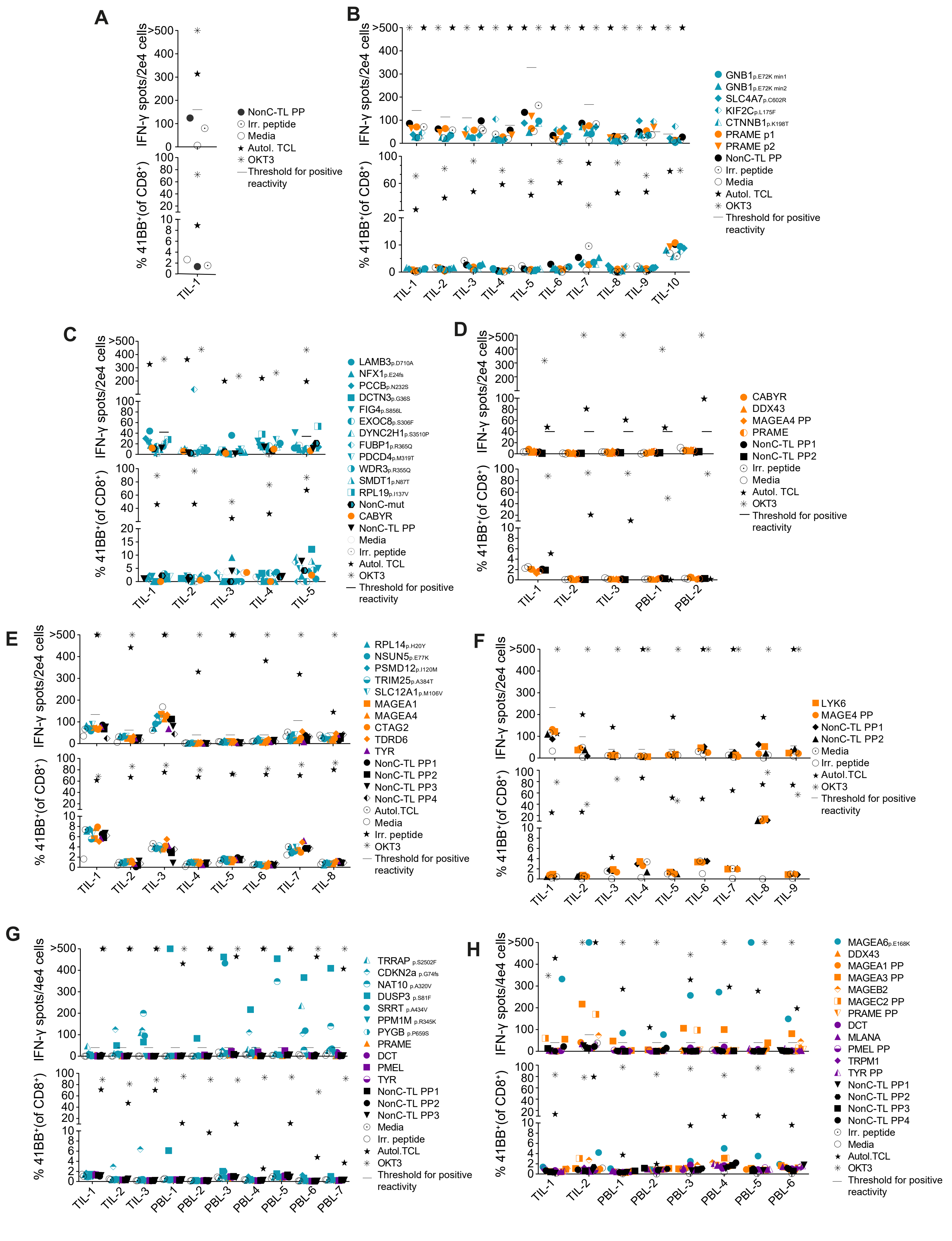
A
IFN-γ spots/2e4 cells
% 41BB+(of CD8+)
TIL-1
NonC-TL PP
Irr. peptide
Media
Autol. TCL
OKT3
Threshold for positive reactivity
B
IFN-γ spots/2e4 cells
% 41BB+(of CD8+)
TIL-1 TIL-2 TIL-3 TIL-4 TIL-5 TIL-6 TIL-7 TIL-8 TIL-9 TIL-10
GNB1 p.E72K min1
GNB1 p.E72K min2
SLC4A7 p.C602R
KIF2C p.L175F
CTNNB1 p.K198T
PRAME p1
PRAME p2
NonC-TL PP
Irr. peptide
Media
Autol. TCL
OKT3
Threshold for positive reactivity
C
IFN-γ spots/2e4 cells
% 41BB+(of CD8+)
TIL-1 TIL-2 TIL-3 TIL-4 TIL-5
LAMB3 p.D710A
NFX1 p.E24fs
PCCB p.N232S
DCTN3 p.G36S
FIG4 p.S856L
EXOC8 p.S306F
DYNC2H1 p.S3510P
FUBP1 p.R365Q
PDCD4 p.M319T
WDR3 p.R355Q
SMDT1 p.N87T
RPL19 p.I137V
NonC-mut
CABYR
NonC-TL PP
Media
Irr. peptide
Autol. TCL
OKT3
Threshold for positive reactivity
D
IFN-γ spots/2e4 cells
% 41BB+(of CD8+)
TIL-1 TIL-2 TIL-3 PBL-1 PBL-2
CABYR
DDX43
MAGEA4 PP
PRAME
NonC-TL PP1
NonC-TL PP2
Irr. peptide
Media
Autol. TCL
OKT3
Threshold for positive reactivity
E
IFN-γ spots/2e4 cells
% 41BB+(of CD8+)
TIL-1 TIL-2 TIL-3 TIL-4 TIL-5 TIL-6 TIL-7 TIL-8
RPL14 p.H20Y
NSUN5 p.E77K
PSMD12 p.I120M
TRIM25 p.A384T
SLC12A1 p.M106V
MAGEA1
MAGEA4
CTAG2
TDRD6
TYR
NonC-TL PP1
NonC-TL PP2
NonC-TL PP3
NonC-TL PP4
Autol.TCL
Media
Irr. peptide
OKT3
Threshold for positive reactivity
F
IFN-γ spots/2e4 cells
% 41BB+(of CD8+)
TIL-1 TIL-2 TIL-3 TIL-4 TIL-5 TIL-6 TIL-7 TIL-8 TIL-9
LYK6
MAGE4 PP
NonC-TL PP1
NonC-TL PP2
Media
Irr. peptide
Autol.TCL
OKT3
Threshold for positive reactivity
G
IFN-γ spots/4e4 cells
% 41BB+(of CD8+)
TIL-1 TIL-2 TIL-3 PBL-1 PBL-2 PBL-3 PBL-4 PBL-5 PBL-6 PBL-7
TRRAP p.S2502F
CDKN2a p.G74fs
NAT10 p.A320V
DUSP3 p.S81F
SRRT p.A434V
PPM1M p.R345K
PYGB p.P659S
PRAME
DCT
PMEL
TYR
NonC-TL PP1
NonC-TL PP2
NonC-TL PP3
Media
Irr. peptide
Autol.TCL
OKT3
Threshold for positive reactivity
H
IFN-γ spots/4e4 cells
% 41BB+(of CD8+)
TIL-1 TIL-2 PBL-1 PBL-2 PBL-3 PBL-4 PBL-5 PBL-6
MAGEA6 p.E168K
DDX43
MAGEA1 PP
MAGEA3 PP
MAGEB2
MAGEC2 PP
PRAME PP
DCT
MLANA
PMEL PP
TRPM1
TYR PP
NonC-TL PP1
NonC-TL PP2
NonC-TL PP3
NonC-TL PP4
Irr. peptide
Media
Autol.TCL
OKT3
Threshold for positive reactivity

**Supplemental Figure 6. Evaluation of pre-existing T-cell responses to candidate tumor antigens in cancer patients.** Reactivity was evaluated by co-incubating from 2e4 to 4e4 T cells (TILs o PBLs sorted based on specific markers, e.g. PD1hi), with 1-2e5 autologous APC with 1 µg/mL of selected peptides either alone or in pools (PP). IFN-γ ELISPOT and 4-1BB upregulation by flow cytometry were used to measure T-cell responses after 20 h. The number of IFN-γ spots per well (top panel) and the percentage of $CD8^+$ T cells expressing 4-1BB (bottom panel) are shown. Mutated peptides are plotted in turquoise, cancer-germline in orange, melanoma-associated in purple, and nonC-TL in black. PP stands for peptide pool. Plotted cells were gated on live $CD3^+CD8^+$ lymphocytes. '>' denotes greater than 500 spots/2e4 cells. B cells or CD4+ were used as APC. Experiments were performed twice. **a** Gyn-1 **b** Gyn-2 **c** Gyn-3 **d** H&N-1 **e** H&N-2 **f** Mel-1 **g** Mel-2. The threshold to consider positive reactivity was calculated as explained in the Methods section and is shown with a line.

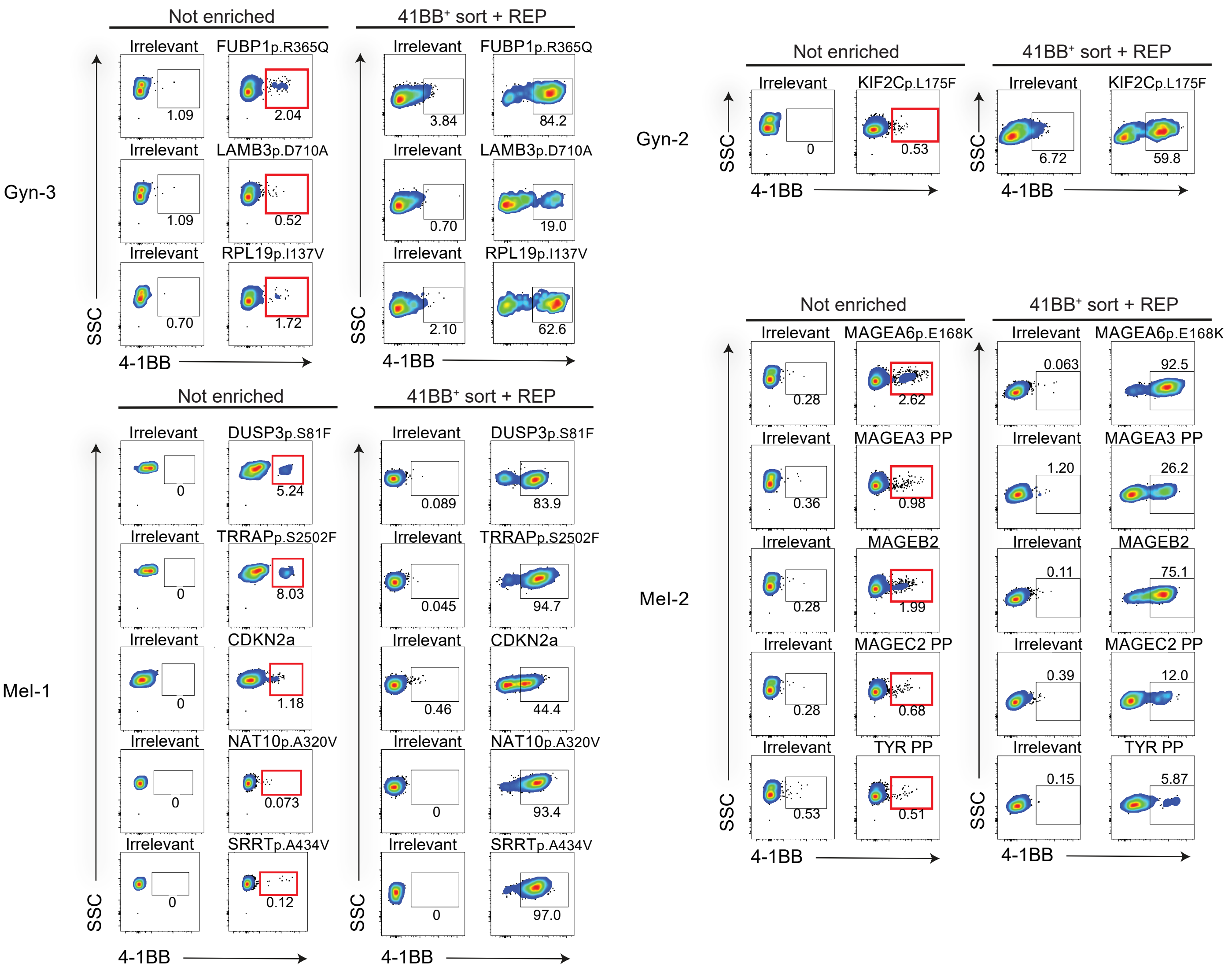


**Supplemental Figure 7. Enrichment of antigen-specific T cells from cancer patients by flow cytometry-based sorting.** Antigen-specific T cells from TILs or PBLs were enriched by flow cytometry sorting of CD3$^+$CD8$^+$4-1BB$^+$ lymphocytes after 20 h co-culture with autologous APC pulsed with the peptides indicated (red squares). Sorted cells were expanded for 14 days using a rapid expansion protocol (REP). To evaluate the enrichment, 2e4 expanded T cells were co-cultured with autologous APC pulsed with the peptides specified and the activation was measured based on 4-1BB upregulation after 20 h by flow cytometry. Dot plots display the percentage of 4-1BB cells after gating on live/CD3$^+$CD8$^+$. Autologous APC pulsed with an irrelevant peptide were used as a negative control.

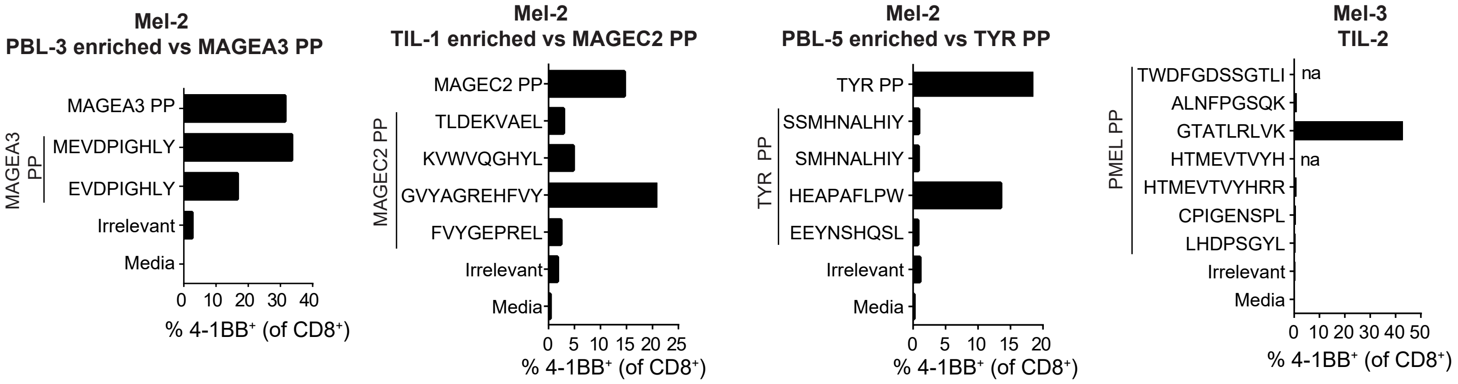


**Supplemental Figure 8. Deconvolution of peptide pools containing recognized CGA or melanoma-associated antigens.** To determine the specific peptide recognized within the peptide pool (PP), 2e5 autologous APC were pulsed with individual peptides and co-cultured with 2e4 reactive lymphocyte populations (with or without previous enrichment by sorting). After 20 h, lymphocyte activation was measured based on 4-1BB upregulation by flow cytometry. Cells were gated on live/CD3⁺CD8⁺ lymphocytes. Autologous APC pulsed with an irrelevant peptide were used as a negative control.

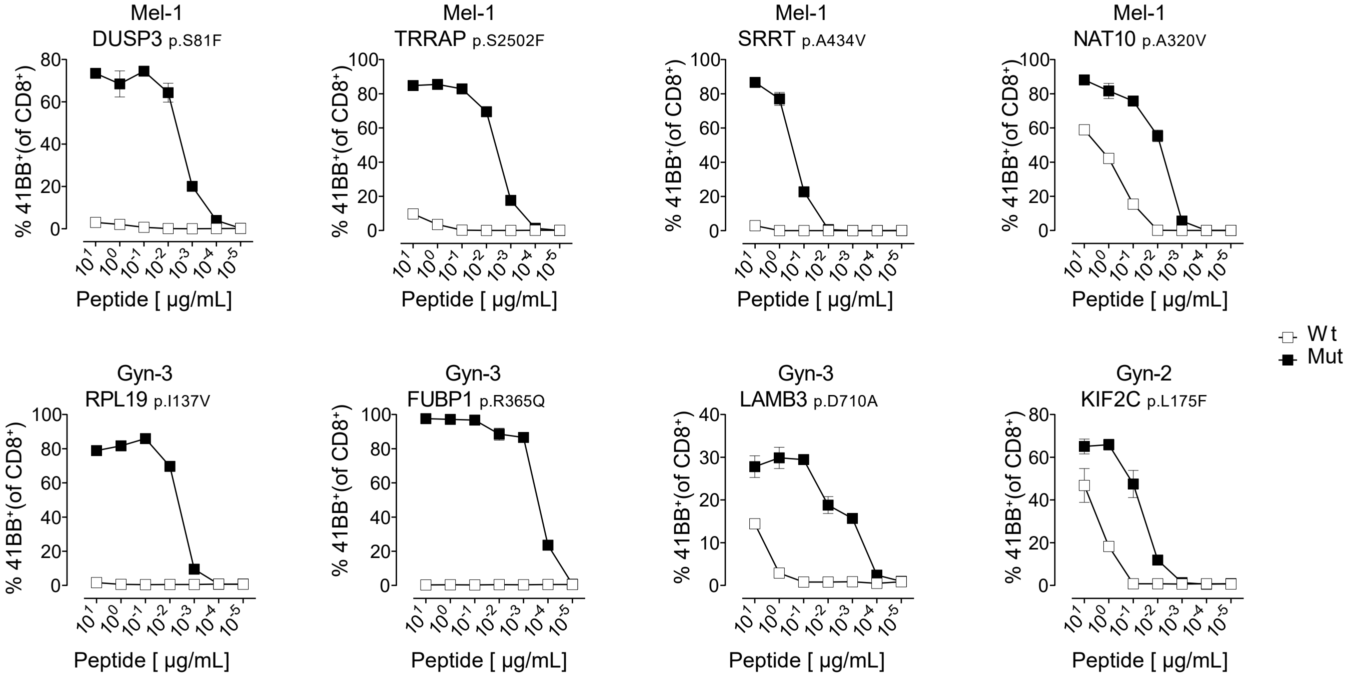


**Supplemental Figure 9. Functional avidity and specificity of neoantigen-specific T cells isolated from cancer patients.** CD8⁺ lymphocyte populations recognizing the mutated HLA-I peptides indicated were enriched by flow cytometry sorting based on 4-1BB expression and expanded for 14 days. To evaluate the avidity and specificity of the enriched antigen-specific populations, 2e4 T cells were incubated with 1-2e5 autologous B cells or CD4+ T cells pulsed with serial dilutions of the wild type (Wt) or mutant HPLC versions. After 20 h, 4-1BB upregulation was measured by flow cytometry. Cells were gated on live/CD3⁺CD8⁺ lymphocytes. Experiments were performed twice with technical duplicates. SD is plotted.

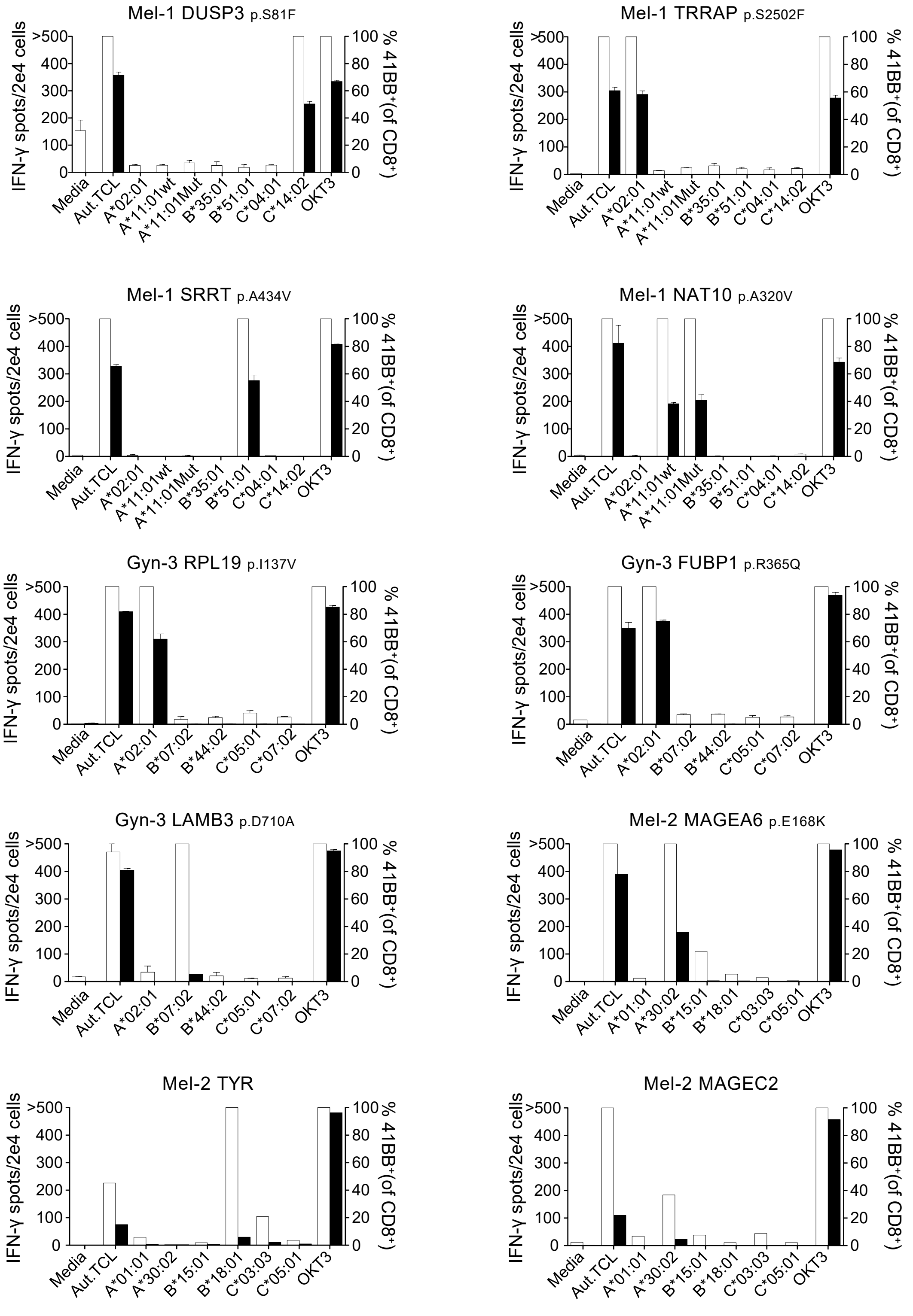


**Supplemental Figure 10. HLA restriction element and recognition of autologous tumor cell lines by antigen-specific T cells isolated from cancer patients.**

$CD8^+$ lymphocyte populations recognizing the HLA-I peptides indicated were enriched by flow cytometry sorting based on 4-1BB expression and expanded for 14 days. The recognition of autologous TCL (Aut.TCL) was evaluated by co-incubating from 3e4 to 5e4 tumor cells with 2e4 T cells. COS7 cells co-transfected with the indicated individual HLA-I alleles followed by peptide pulsing were used as target cells to determine the HLA restriction element. Tumor cell and COS7 recognition was evaluated by IFN-γ release (left axis, white bars) and $4\text{-}1BB^+$ upregulation (right axis, black bars). '>' denotes greater than 500 spots/2e4 cells. OKT3 and media were used as positive and negative controls, respectively. Experiments were performed twice with technical duplicates. SD is plotted.

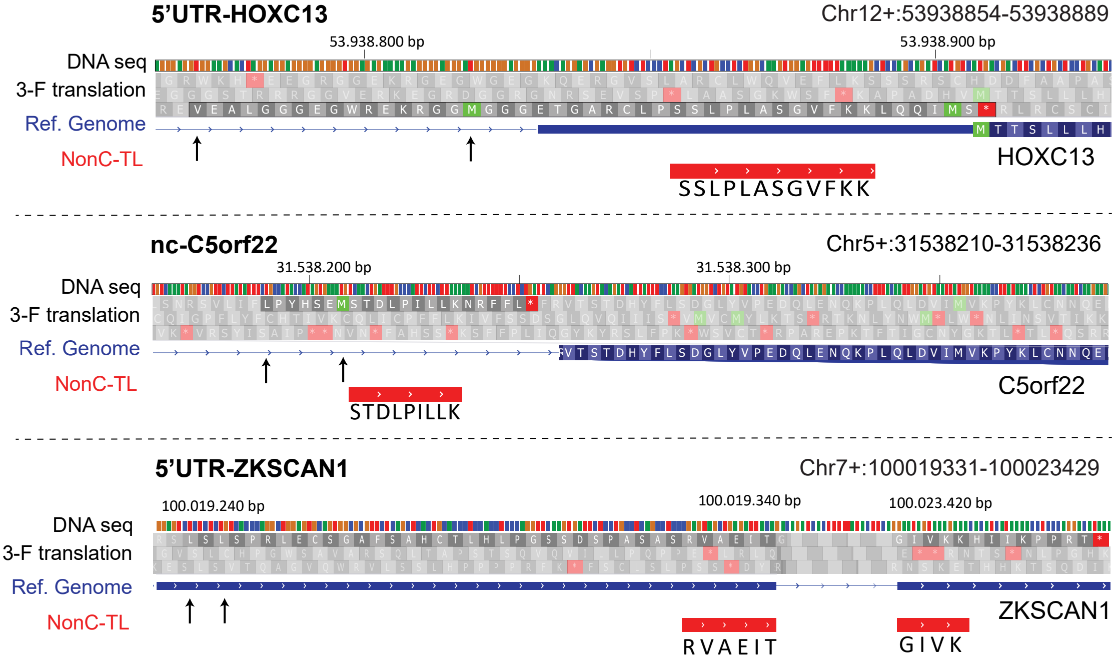


**Supplemental Figure 11. Genomic location of the three immunogenic nonC-TL identified through IVS.** Annotated translated sequences are depicted in blue and the nonC-TL sequence of interest are shown in red. The potential start codons are marked with an arrow. The predicted ORF is highlighted. Image modified from IGV viewer.

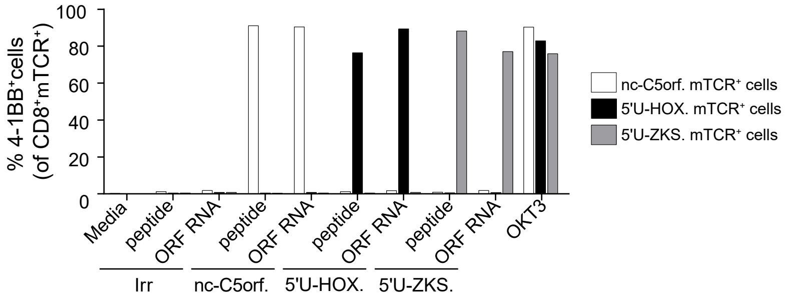


**Supplemental Figure 12. Recognition of the predicted ORFs encoding the immunogenic nonC-TL by TCR transduced cells.** The DNA sequences of the predicted open reading frames (ORF) for each immunogenic nonC-TL were cloned into pcDNA3.1 without any modification except for a Kozak sequence upstream, and the RNA was generated by in vitro transcription. PBL transduced with the antigen-specific TCR specified were co-cultured with B cells either electroporated with the RNA encoding the predicted or an irrelevant ORF or pulsed with the nonC-TL or an irrelevant peptide. After 20 h, T-cell activation was evaluated by measuring 4-1BB upregulation by flow-cytometry of live CD8$^{+}$mTCRtd$^{+}$ cells.

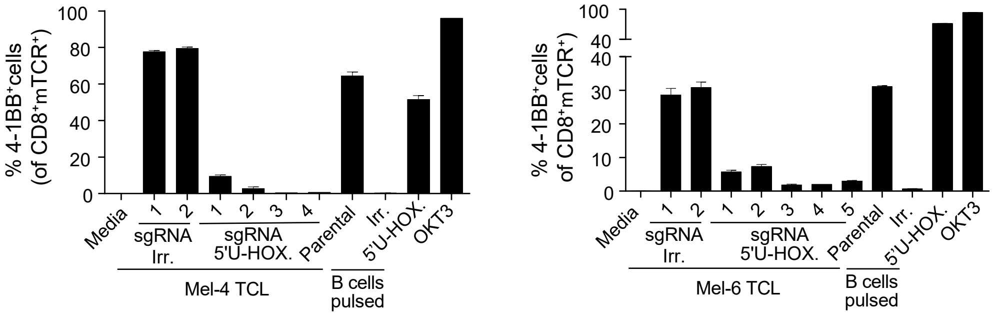


**Supplemental Figure 13. Loss of recognition of TCL transfected with Cas9-sgRNA targeting the predicted 5'U-HOXC13 ORF.** Patient-derived tumor cell lines Mel-4 (right) and Mel-6 (left) were transfected with lentiviruses encoding for Cas9 together with an irrelevant sgRNA or different sgRNA specifically targeting the genomic location of the immunogenic nonC-TL 5'U-HOXC13. Following puromycin selection, TCL were co-cultured with PBL transduced with the 5'U-HOXC13-specific TCR and T-cell activation was measured after 20 h by flow cytometry. Plotted cells were gated on live CD8$^{+}$mTCR$^{+}$ T cells. The experiment was performed with technical duplicates, mean and SD are plotted.

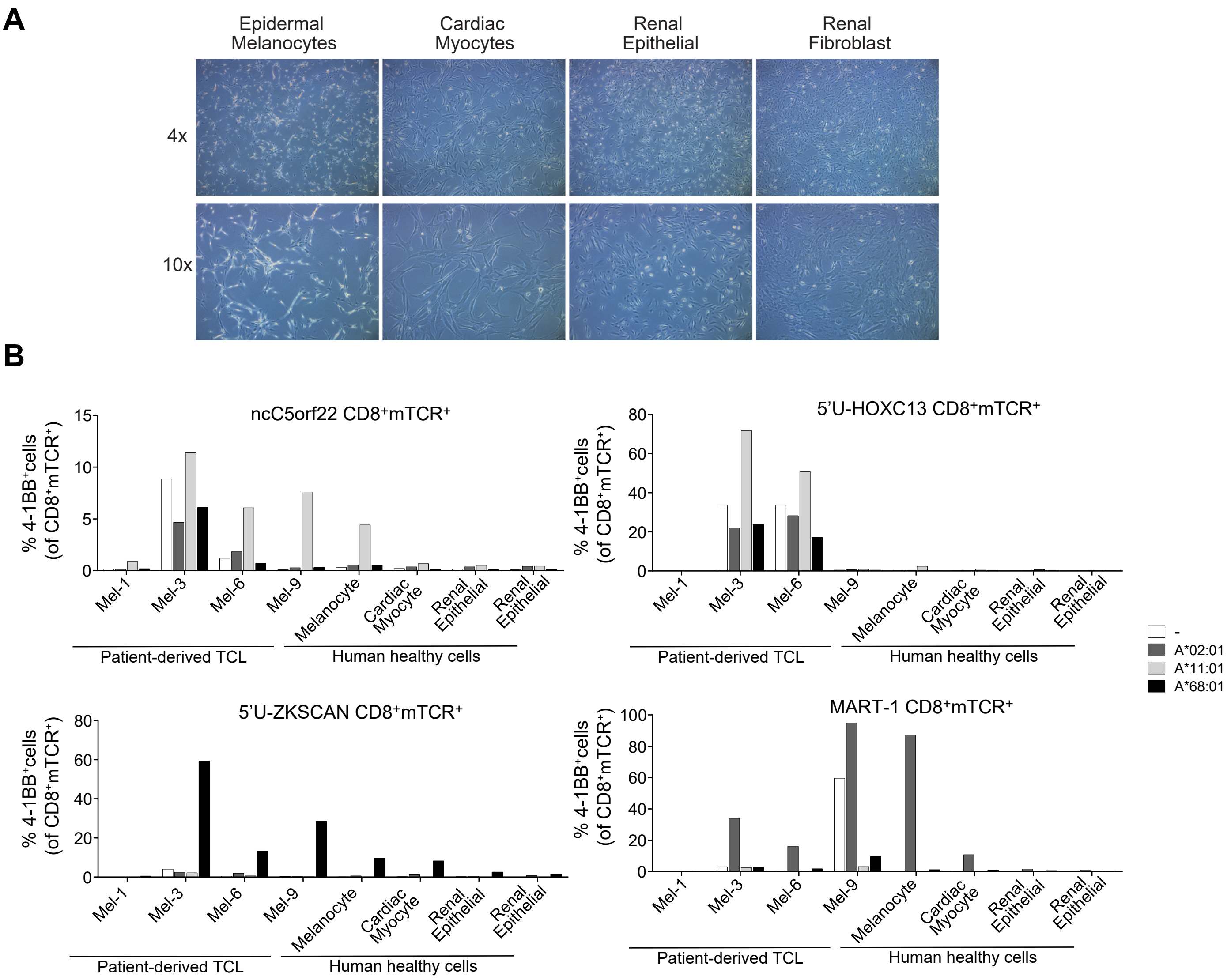


**Supplemental Figure 14. Assessment of expression and translation of the immunogenic nonC-TL in human healthy cells.** **a** Microscopy images of epidermal melanocytes, cardiac myocytes, renal epithelial cells, and renal fibroblast from healthy donors. **b** The expression and translation of the nonC antigens was empirically evaluated by co-culturing $CD8^+$ cells transduced with TCR recognizing the antigens specified (CD8+TCRtd+) with human healthy cells of different origin and previously tested melanoma TCLs electroporated with the HLA alleles specified (x-axis). After 20h, the lymphocyte activation was evaluated by measuring 4-1BB upregulation by flow-cytometry (y-axis). Plotted cells are gated on live/$CD3^+CD8^+$/$mTCR^+$.

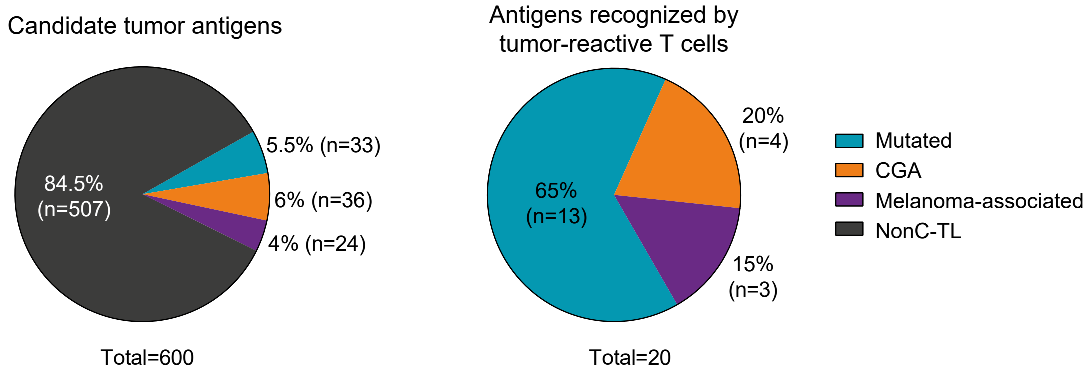


**Supplemental Figure 15. NonC-TL are the main source of candidate tumor antigens identified by proteogenomics but tumor-reactive T cells preferentially recognize neoantigens derived from NSM.**

The percentage and absolute number of candidate tumor antigens presented on HLA-I identified through proteogenomics in the 9 patient-derived TCL and tested for pre-existing T cell responses, are depicted by antigen category (left). The absolute number and percentage of HLA-I ligands recognized by tumor-reactive lymphocytes isolated from cancer patients are depicted by antigen category (right).

**Supplemental Table 1. Patient characteristics and relevant experimental information**

| Patient ID | Tumor type | HLA-I typing[a] | #Cells/IP | #IP | #Runs | #NSM[f] | #Mutated HLA-I ligands[g] |
|---|---|---|---|---|---|---|---|
| **Gyn-1** | Endometrial cancer, CNL | A*03:01<br>B*47:01 B*51:01<br>C*02:02 C*06:02 | 5e7 | 2 | 4 | 110 | 0 |
| **Gyn-2** | Endometrial cancer, POLE | A*02:01<br>B*51:01<br>C*02:02 C*15:02 | 1e8 | 1 | 2 | 14765 | 13 |
| **Gyn-3** | Endometrial cancer, POLE | A*02:01[b]<br>B*07:02[c] B*44:02[c]<br>C*05:01 C*07:02 | unk. | 2 | 4 | 7512 | 5 |
| **Gyn-4** | Ovary small cell carcinoma | A*01:01 A*02:01<br>B*27:05 B*51:01<br>C*01:02 C*15:02 | 1.2e8 | 2 | 4 | 14 | 0 |
| **H&N-1** | Hypopharyngeal carcinoma | A*02:01[d] A*03:01[b]<br>B*07:02[b] B*18:01[e]<br>C*05:01 C*07:02 | 2e8 | 3 | 6 | 2485 | 5 |
| **H&N-2** | Oropharyngeal squamous cell carcinoma | A*24:02 A*29:02<br>B*14:02 B*44:03<br>C*02:02 C*16:01 | 6e7 | 2 | 4 | 266 | 0 |
| ***Mel-1** | Melanoma | A*02:01 A*11:01[b]<br>B*35:01 B*51:01<br>C*04:01 C*14:02 | 5.5e7 | 2 | 4 | 275 | 3 |
| ***Mel-2** | Melanoma | A*01:01 A*30:02<br>B*15:01 B*18:01<br>C*03:03 C*05:01 | 6e7-9e7 | 3 | 6 | 255 | 1 |
| **Mel-3** | Melanoma | A*11:01 A*29:02<br>B*35:01 B*35:02<br>C*04:01 | 1e8-1.4e8 | 3 | 8 | 2951 | 7 |

[a]HLA-I typing was evaluated for TCL and matched PBMC WES data. Specific mutations of unknown functional impact detected in the HLA molecules of the TCLs are noted. [b]Missense mutations. [c]Missense in one of the HLA-B alleles. [d]Loss of heterozigosity. [e]Nonsense mutations. [f]Total number of non-synonymous mutations NSM identified by WES. CNL; copy number low, POLE; DNA Polymerase Epsilon, Catalytic Subunit, IP; Immunoprecipitation. [g]HLA-I ligands derived from NSM identified by proteogenomics.
